# Urine single cell RNA-sequencing in focal segmental glomerulosclerosis reveals inflammatory signatures in immune cells and podocytes

**DOI:** 10.1101/2020.10.18.343285

**Authors:** Khun Zaw Latt, Jurgen Heymann, Joseph H. Jessee, Avi Z. Rosenberg, Celine C. Berthier, Sean Eddy, Teruhiko Yoshida, Yongmei Zhao, Vicky Chen, George W. Nelson, Margaret Cam, Parimal Kumar, Monika Mehta, Michael C. Kelly, Matthias Kretzler, The Nephrotic Syndrome Study Network (NEPTUNE), The Accelerating Medicines Partnership in Rheumatoid Arthritis and Systemic Lupus Erythematosus (AMP RA/SLE) consortium, Cheryl, A. Winkler, Jeffrey B. Kopp

## Abstract

The diagnosis of focal segmental glomerulosclerosis (FSGS) requires a renal biopsy, which is invasive and can be problematic in children and in some adults. We used single cell RNA-sequencing to explore disease-related cellular signatures in 23 urine samples from 12 FSGS subjects. We identified immune cells, predominantly monocytes, and renal epithelial cells, including podocytes. Analysis revealed M1 and M2 monocyte subsets, and podocytes showing high expression of genes for epithelial-to-mesenchymal transition (EMT). We confirmed M1 and M2 gene signatures using published monocyte/macrophage data from lupus nephritis and cancer. Using renal transcriptomic data from the Nephrotic Syndrome Study Network (NEPTUNE), we found that urine cell immune and EMT signature genes showed higher expression in FSGS biopsies compared to minimal change disease biopsies. These results suggest that urine cell profiling may serve as a diagnostic and prognostic tool in nephrotic syndrome and aid in identifying novel biomarkers and developing personalized therapeutic strategies.

## Introduction

Focal segmental glomerulosclerosis (FSGS) and minimal change disease (MCD) are the major causes of nephrotic syndrome with similar clinical features. It is therefore important to arrive at the correct diagnosis between these two diseases in order to initiate the effective treatment. MCD patients usually respond well to steroid therapy and usually have excellent long-term prognosis. In contrast, FSGS patients are typically resistant to steroid therapy and show progressive loss of glomerular filtration rate (GFR) ^1, 2^. Currently, renal biopsy is the principal method for the histological diagnosis of nephrotic syndrome. However, it is an invasive procedure that is sometimes deferred, particularly in children, and is typically performed only once in adults. Due to sampling limitations and the focal distribution of lesions throughout the renal parenchyma in FSGS, biopsy can also fail to distinguish MCD from early FSGS. Moreover, the current approaches to renal biopsy analysis provide limited information about molecular mechanisms of complex diseases, such as FSGS.

In recent years, single cell RNA sequencing (scRNA-seq) has emerged as a powerful tool to characterize single cell transcriptomes from various sources. Several reviews of this methodology, applied to kidney research, have been published recently ^3, 4, 5, 6^. However, these studies applied single cell or nuclear RNA-seq approach directly to kidney tissue. We hypothesized that urine from patients with kidney diseases could be a useful, non-invasive source of information about the disease and urine scRNA-seq could add valuable transcriptional information about injured primary renal parenchymal cells that appear in the urine, including podocytes and tubular epithelial cells as well as reactive cells such as immune cells, and could distinguish these cells from urothelial cells. In order to evaluate the urine cells for potential application of urine as a diagnostic tool for FSGS and to uncover the molecular mechanisms of the disease at the single cell level, we performed scRNA-seq of urine samples from subjects with FSGS.

## Results

### Study design for single cell RNA sequencing of FSGS urine cells

To parse the gene expression signatures of single cells in the urine, we performed scRNA-seq of 23 urine cell samples from 12 FSGS subjects recruited at the National Institutes of Health Clinical Center, including seven samples collected longitudinally from one subject. Using the known canonical marker gene sets, we identified podocytes, tubular epithelial cells and immune cells (monocytes and lymphocytes). We found that these marker genes were also highly expressed in transcriptomic data obtained from kidney tissue by evaluating expression levels of these genes in glomerular and tubulointerstitial expression data from the Nephrotic Syndrome Study Network (NEPTUNE) cohort. We were able to distinguish FSGS and MCD subjects based on the expression of these genes. The majority of immune cells in the urine samples were monocytes and among these we identified M1 and M2 monocyte subtypes with distinct gene expression profiles. We used *in silico* approaches to annotate M1 and M2 subtypes and to show that their gene expression signature was similar across several inflammatory conditions.

### Identification of different cell types in the urine of FSGS patients

We used Seurat package ^7^ for data analysis, including count normalization, cell filtering and cell clustering and we used the Harmony package ^8^ to correct for batch effects across samples. These analyses produced 15 cell clusters (**Fig. 1A**), with cells from multiple samples in most clusters (**Fig. 1B, Fig. S1 and Table S1**). We used known canonical marker genes for kidney and immune cells to identify the cell type for each cluster (**Fig. S2**).

**Fig. 1.**
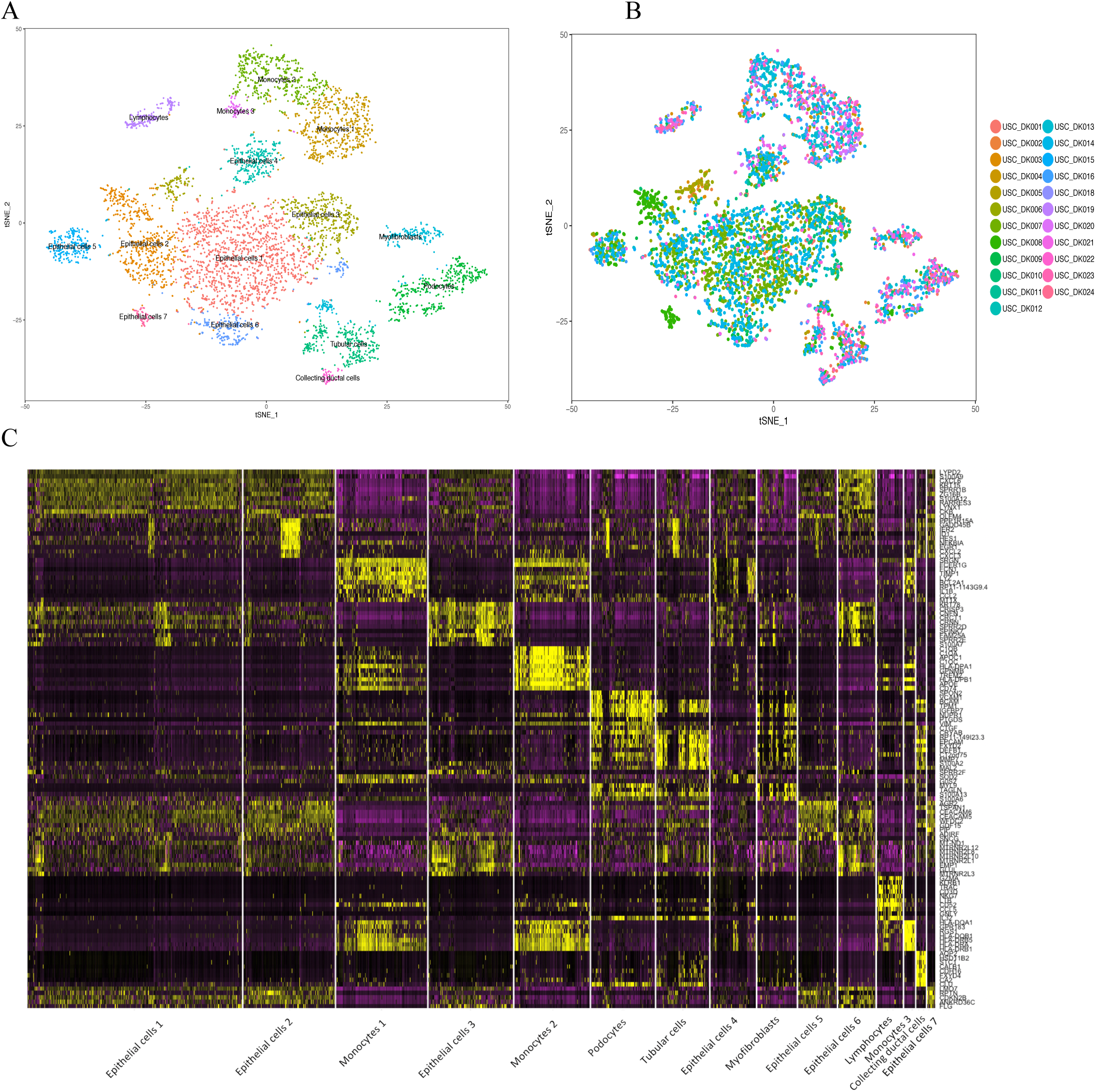
As shown in t-SNE plots, gene expression profiles of human urine cells indicate four distinct categories of cells (**A**) Batch-corrected t-SNE plot of urine single cell aggregate data from all 23 urine samples, showing 14 individual cell clusters and cell types. (**B**) Shown is the same t-SNE plot with a different color for each sample. (**C**) Principle component heatmap plot showing ten most highly expressed genes in each of 14 clusters (vertical columns), with each row representing one gene, with high expression (yellow), intermediate expression (purple) and low expression (black).

We identified a podocyte cluster that expressed *WT1, PLA2R1* and *SYNPO* (**Fig. S2C**) and also for the parietal epithelial cell (PEC) markers, such as *PAX2, PAX8* and *CLDN1* (**Fig. S3**). Other canonical podocyte marker genes such as *PODXL* and *NPHS1* were not strongly expressed in the urinary podocytes, which may reflect an altered transcriptional state of podocytes that were shed into the urine. We compared the gene expression profile of the podocyte cluster with all the remaining cell clusters. We found that for the podocyte cluster, the most highly expressed gene with the highest logarithmic (log)- fold change was *IGFBP7* (2.55 fold), a marker that had also been reported for podocytes ^9^. Among other top expressed genes were myofibroblast markers (*CTGF, MYL9*), mesenchymal markers (*VIM, THY1*), extracellular matrix proteins (*MMP7, CAV1*), markers for smooth muscle differentiation (*CALD1, TPM1* and *TAGLN*) and *CRYAB*, which induces epithelial-to-mesenchymal transition (EMT) ^10^ (**Table S2**).

Highly expressed genes distinctive for the tubular cell cluster included *POU3F3, UMOD*, and *FXYD2*. This cluster also showed high expression of *TPM1* and *CRYAB*, consistent with EMT. There were also a small collecting ductal cell cluster, identified by high expression of *AQP2*, and a myofibroblast cluster, identified by high expression of *TAGLN, MYL9* and *ACTA2*. The remaining clusters were seven epithelial cell clusters positive for *KRT6A, KRT13, KRT15* and *KRT17*, and likely originated from different segments of the male and female genitourinary tracts (**Fig. S2**).

We identified three monocyte clusters (MC1, MC2 and MC3), comprising a total of 1040 cells, which were positive for *CD14* and *FCGR3A* (CD16). The three monocyte clusters shared some highly expressed genes, including *FCER1G, TYROBP* and *HLA* genes. The most highly expressed genes for MC1 included *TIMP1, CCL2* and *IL1B*. The most highly expressed genes for MC2 included *APOE, C1QB* and *APOC1*. MC3 showed strong expression of *HLA* class II genes and *CD74* (**Fig. 1C**), suggesting the differentiation towards dendritic cells and manifesting active antigen presentation to T cells ^11^. This cluster also had the highest expression of dendritic cell marker genes such as *CD1C, CD1E, CCR7, FCER1A* and *CLEC10A* (**Fig. S4**). We observed a single lymphocyte cluster expressing *CD3G* and *GZMA*. The lymphocyte cluster was mostly composed of T lymphocytes with high expression of cytotoxic genes, including *GNLY, GZMA* and *LTB*, and a smaller subgroup of B cells expressing *CD19* and *MS4A1* (**Fig. 1C and Fig. S2A)**.

The Harmony program returns batch-corrected principal components but does not return corrected gene expression levels. We reasoned that corrected expression levels could be approximated by reversing the singular value decomposition (SVD) that generates the “principal components” (technically embeddings from the SVD, but commonly referred to as PCs). Thus, we calculated corrected expression levels by reversing the SVD using the Harmony-corrected PCs and the uncorrected embeddings and singular values. Here we worked from the ansatz that the Harmony correction moves the cell embeddings in PC space, but does not significant change the SVD, thus the SVD can be approximately reversed using the original loadings matrix and singular values. We demonstrate that these back-calculated gene expression levels embody the Harmony corrections to good approximation by redoing the SVD with these expression levels, and showing that resulting cell embeddings, without correction, substantially reproduce the Harmony batch corrections **(Fig. S5 and S6)**.

**Fig S5. A, B**, and **C** compare PCs 1 and 2 for uncorrected, Harmony corrected, and uncorrected PCs from the SVD on the back-calculated expression levels; B and C are nearly identical. **Fig. S6** shows tSNE plots colored by back-calculated cell expression levels, for comparison with cell expression level plots in **Fig. S1**; the distribution of genes over the clusters is very similar.

### Pathway Analysis

We performed gene ontology (GO) pathway analysis for each cell cluster, selecting significantly expressed genes, defined as those with an adjusted p value < 0.05 when compared with all remaining cells. Notably, the podocyte cluster showed the highest number of significantly expressed genes (n=1492). Therefore, we selected the most highly expressed genes with an average log-fold change more than 1 for the pathway analysis (n = 38) (**Table S2**). The significantly activated pathways included the pathways for cell adhesion, which is likely important for interaction with other cell types and with extracellular matrix; muscle organ development, which may reflect EMT; and extracellular matrix organization, wound healing and collagen biosynthesis processes (**Table S3**).

The most significant up-regulated pathways for the monocyte and lymphocyte clusters included signal recognition-dependent co-translational protein targeting to membranes and translational initiation. Both of these processes are consistent with the active secretion of molecules, possibly cytokines and signaling molecules. The three monocyte clusters showed enrichment of genes involved in immune processes, such as antigen processing and presentation and interferon-gamma-mediated signaling pathway (**Tables S4 and S5**). Similarly, the lymphocyte cluster manifested enrichment of gene expression in pathways related to T-cell mediated cytotoxicity (**Tables S6 and S7**), consistent with the T cell predominance in that cluster.

### Comparison of RNA expression profiles from FSGS monocytes with healthy PBMC monocytes identified M1 and M2 subpopulations

Since monocytes were the major immune cell type in the FSGS urine, as shown above, we analyzed the monocytes by comparing all three urine monocyte clusters with peripheral blood monocyte data available from 10x Genomics (which was generated using PBMC 8k v2 chemistry) (https://support.10xgenomics.com/single-cell-gene-expression/datasets/2.1.0/pbmc8k?). We found that the most highly expressed genes in both the MC1 and MC2 were upregulated when compared with PBMC monocytes, in which many of these genes were expressed in only a small fraction of cells (**Table 1 and Fig. 2A**).

**Table 1.**
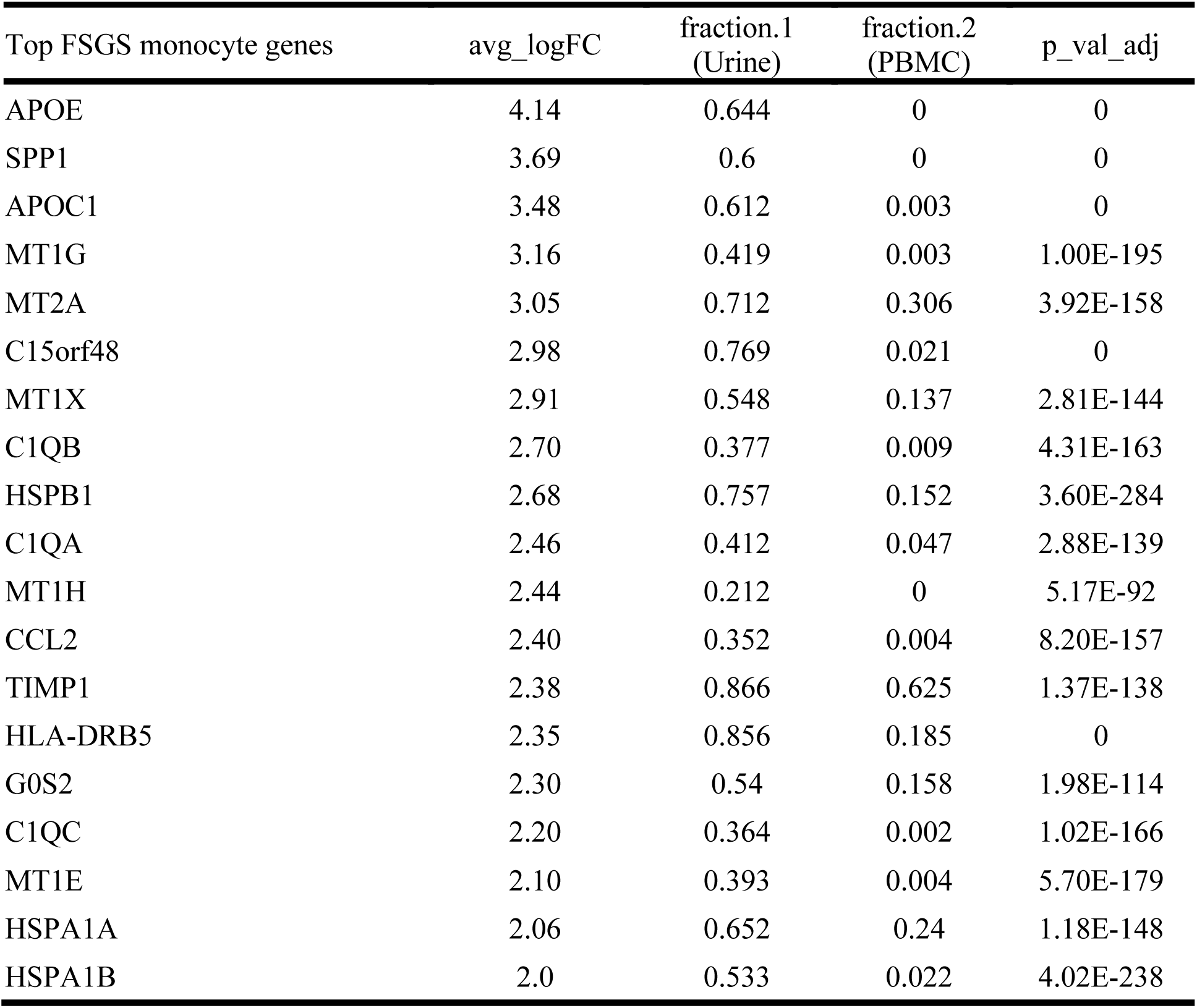
Shown are the top up-regulated genes in FSGS monocytes from all 23 urine samples (n = 1040 cells) compared with healthy peripheral blood monocytes from one healthy subject (n = 1906 cells). The latter data are from 10x Genomics PBMC version 2 with ∼ 8000 cells. Genes are ordered in descending expression levels, shown as average log-fold change (natural log) compared to healthy blood monocytes. Fraction.1 and fraction.2 are the fractions of monocytes in urine and peripheral blood, respectively, that express mRNA for these genes.

**Fig. 2.**
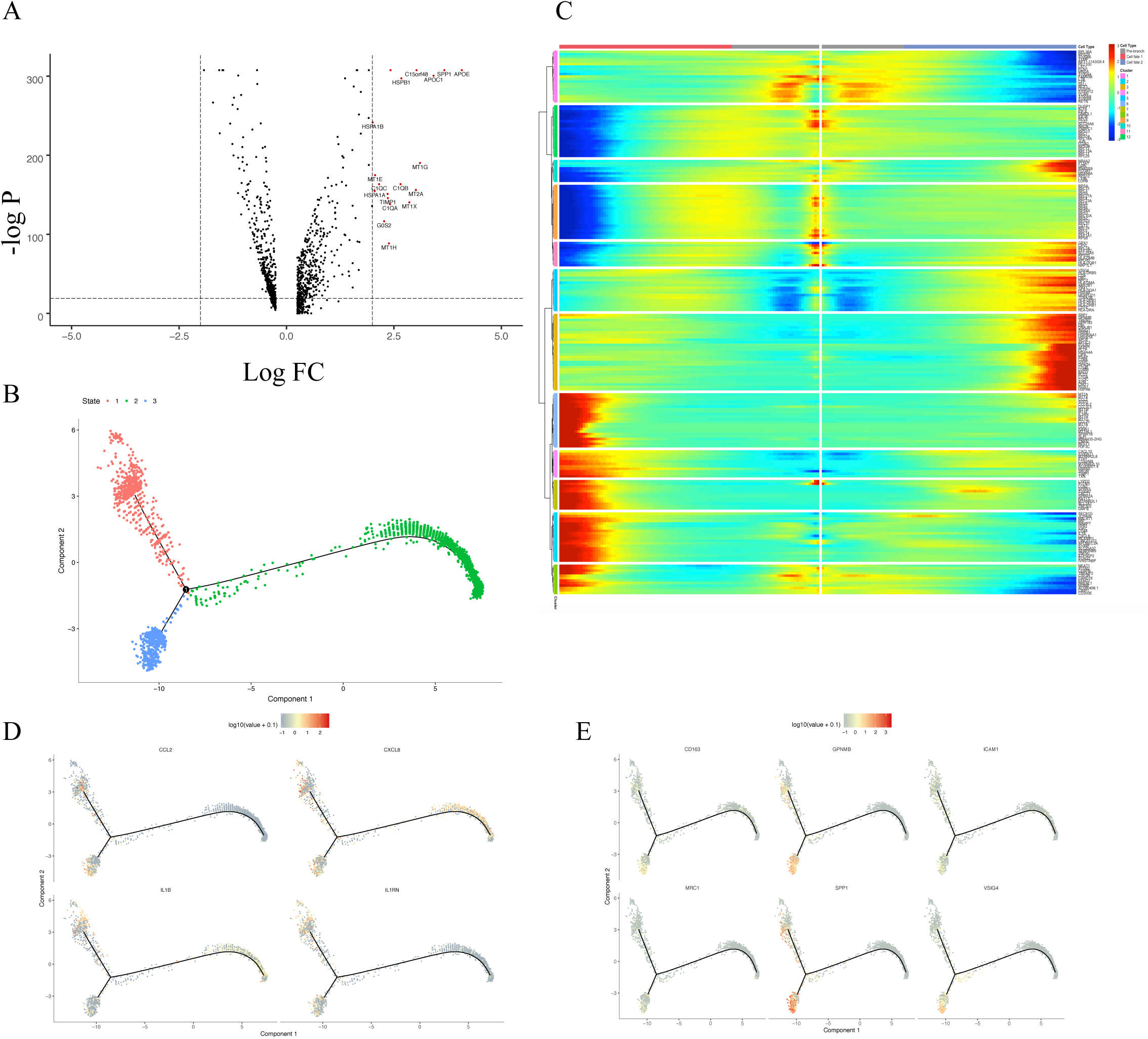
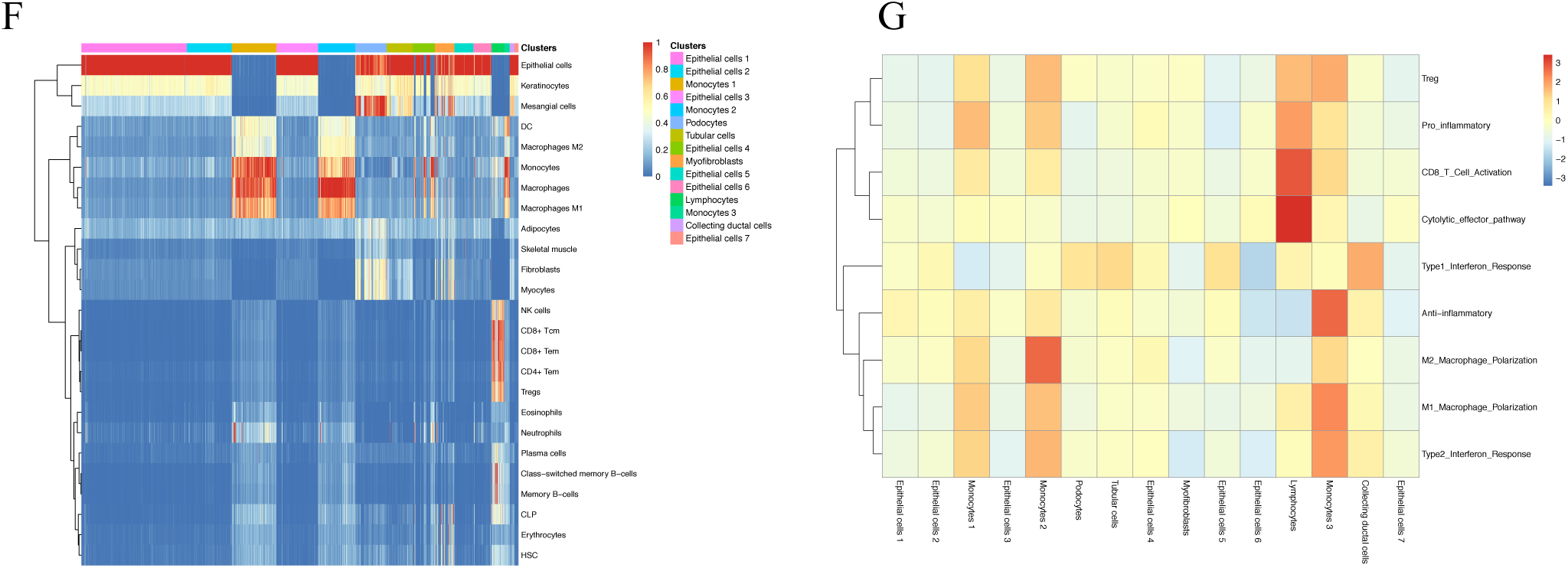
(**A**) Volcano plot showing the up- and down-regulated genes in FSGS monocytes compared with healthy PBMC monocytes (red dots show genes with log fold change > 2). (**B**) Pseudotime trajectory analysis of combined FSGS and peripheral blood monocytes. Trajectory analysis shows a branching point, connecting three states. Healthy peripheral blood monocytes are all in state 2 (green) as shown in **Fig. S7**. (**C**) Expression heatmap of these three states (branches), considering peripheral blood monocytes as the naïve state (pre-branch). Each horizontal line represents one gene and the vertical lines represent all 2946 monocytes (1040 from urine of 12 FSGS subjects and 1904 from peripheral blood of one healthy donor). Expression level is color coded, from red (high) to blue (low). Only genes with p-values less than 10^−20^ are shown here. (**D**) Shown is expression of canonical M1 monocyte marker genes in pseudo-time branches. (**E**) Shown is expression of canonical M2 monocyte marker genes in pseudo time branches. (**F**) Urine single cell data was annonated by SingleR R-package using Blueprint and Encode reference data (transcriptional data for various cell types). In this matrix, each vertical line represents one urine cell and the horizontal lines represent comparisons to signatures of 25 most closely matched cell types, as labelled on the right. Blue denotes low enrichment and red denotes high enrichment for the characteristic cell signature as labelled. (**G**) We looked for enrichment of immune signatures using gene lists from Azizi et al. X-axis shows 14 cell clusters and Y-axis shows 9 immune functions. Collective gene signatures are shown as colors representing relative expressions with red color representing highest expression. The monocyte 1 cluster is modestly enriched for M1 polarization and pro-inflammatory gene expression. The monocyte 2 cluster is strongly enriched for M2 polarization and the monocyte 3 cluster is enriched for M1 polarization, and surprisingly, anti-inflammatory pathways. The lymphocyte cluster is enriched for CD8 T cell activation and cytolytic effector pathway genes.

To further characterize the subpopulations of these monocytes and their activated states, we performed pseudotime analysis in Monocle2 (version 2.10.1) ^12^. We combined FSGS monocytes and PBMC monocytes and the analysis gave three branches of cells (**Fig. 2B**), with all PBMC monocytes concentrated at the terminal of branch 2 and FSGS monocytes diverging into branch 1 and 3 (**Fig. S7**). We considered the PBMC monocytes as being in the naïve state (root) and evaluated the significant gene expression changes in pseudotime. Our analysis showed that the FSGS monocytes in branch 1 manifested upregulation of the top genes in MC1 (*TIMP1, CCL2, IL1B*) and those in branch 3 displayed upregulation of the top genes in MC2 (*APOE, APOC1, C1QB*) (**Fig. 2C**).

As the monocytes in branch 1 express *IL1B* and *CCL2*, which are characteristic M1 genes, and those in branch 3 express characteristic M2 genes such as *CD163, MRC1* and *VSIG4* (**Fig. 2D and 2E**), we hypothesized that these two branches represent M1 and M2 populations. We undertook computational approaches to test the hypothesis. We annotated the urine single cell data with Blueprint and Encode reference data using SingleR R package ^13^. Using these datasets, we found that the MC1 expression was more typical for an M1 signature and the MC2 expression was more typical for an M2 signature (**Fig. 2F**). We also evaluated the enrichment of different immune signatures using the gene lists from Azizi et al ^14^, and we found that MC2 showed the strongest enrichment for an M2 signature (**Fig. 2G**). Consistent with the positivity of dendritic cell marker genes, MC3 showed dendritic cell signature enrichment with Blueprint and Encode annotation and strong enrichment of M1 and anti-inflammatory immune signatures (**Fig. 2F and 2G**), suggesting that monocytes in these clusters were differentiating into inflammatory dendritic cells.

To confirm our analyses and to evaluate how generalizable these gene signatures were across different inflammatory conditions, we evaluated publicly available single cell data derived from macrophages from untreated melanoma ^15^ and head and neck cancer ^16^, available at the VirtualCytometry website ^17^ (https://www.grnpedia.org/cytometry) and human kidney allograft rejection data from Humphreys Laboratory website (http://humphreyslab.com) ^18^. In these datasets, we used *IL1B* expression as the marker for M1 macrophages and *APOE* expression for M2 macrophages. We found that across all three conditions listed above, there was a remarkable separation of M1 from M2 macrophages, based on the expression of these two genes (**Fig. S8 and S9**). In melanoma and head and neck cancer, the correlation analysis showed that there were very few cells which had high expression of both genes (**Fig. S8C and S8F)**. We selected *APOE*^+^ macrophages from these cancers and compared differential gene expression with the remaining macrophages. We found that many of the genes with highest log-fold changes overlapped with the top M2 genes from our study, further supporting identification of these genes as a gene set typical of M2 macrophage (**Tables S8 and S9**). For tumor M1 macrophages, gene expression profiles were less similar to FSGS urine M1 monocytes compared to tumor M2 macrophages and urine M2 monocytes, although the former shared expression of some of the most highly expressed genes. (**Tables S10 and S11**).

We also examined the expression of *PLAUR*, which encodes suPAR, a circulating factor implicated in FSGS pathogenesis ^19^, in our urine single cell data. *PLAUR* expression was the highest in the monocyte clusters, compared to other urine cell clusters. We also evaluated *PLAUR* expression in the single cell data from the Humphreys Laboratory. There, we observed that *PLAUR* was highly expressed only in the monocyte clusters found in kidney allograft rejection data ^18^ and, in contrast, relatively low expression was observed in the normal kidney tissue ^20^ (**Fig. S10**). Together, these data suggest that in these instances, kidney inflammation is characterized by increased monocyte *PLAUR* expression.

### Cell-to-cell Interactions

To identify potential cell-to-cell interactions occurring among the various immune and epithelial cell types in the glomerular and tubulointerstitial microenvironment, we used CellPhoneDB ^21, 22^ (www.cellphonedb.org) which makes statistical inferences based on expression of ligands and the cognate receptors. There were potential interactions between immune and renal epithelial cells involving cytokines from TNF family, *TGFB1* and *IL1B* signaling (**Fig. 3A**), and between lymphocytes and monocytes involving *CCL5* (**Fig. 3B**) which is a chemokine secreted by cytotoxic lymphocytes and plays an active role in recruiting leukocytes into inflammatory sites.

**Fig. 3.**
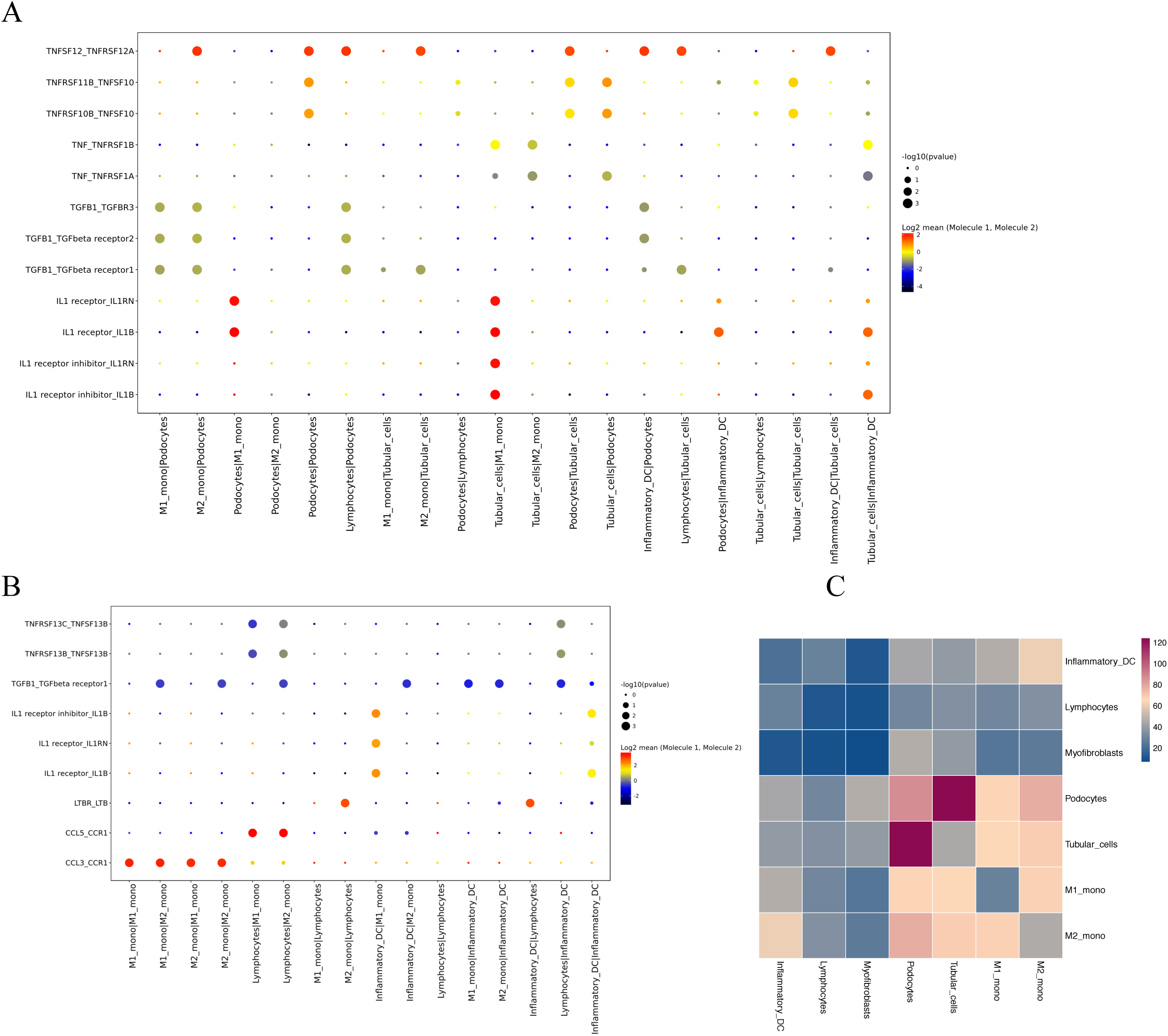
Cell-cell interactions of ligands and receptors between different clusters of urine cells. (**A**) Dot plot showing selected interactions between immune and renal epithelial cell clusters (**B**) Dot plot showing selected interactions between immune cell clusters (**C**) Heat map showing the number of all the interactions between the urine single cell clusters.

Notably, of the TNF family cytokines, the most prominent was the potential *TNFSF12* – *TNFRSF12A* (TWEAK/Fn14) interaction between the immune cells and kidney epithelial cells. Other potential interactions were *TNFSF10* (TRAIL) with *TNFRSF10B* (DR5) and with *TNFRSF11B* (osteoprotegerin) receptors, and *TNF* with *TNFRSF1A* and *TNFRSF1B* receptors.

### Immune and EMT gene signature differences between FSGS and MCD

Since we detected immune cells in the FSGS urine, we hypothesized that the infiltration of these immune cells into the kidney may be important for the development and progression of FSGS. Consequently, we sought to evaluate whether the presence of these immune cells would allow to distinguish FSGS from MCD. To this end, we selected the 16 most highly expressed genes identified in the immune cells (8 genes from monocytes and 8 genes from lymphocytes), based on their log-fold changes and being selectively expressed in these cell types (**Fig. S11 and Table S12**). We also confirmed the specificity of these genes for immune cells in the single nuclear RNAseq data from human adult kidney tissue, as reported by Menon et al. ^23^ and found that the expression of these genes was relatively selective and reached the highest levels in immune cells (**Fig. S12**).

We compared expression of these 16 highly-expressed genes between FSGS and MCD cases in kidney transcriptomic data from the NEPTUNE study ^24^. Compared to MCD, FSGS samples had higher expression levels of these 16 genes and the difference was more profound in the tubulointerstitial compartment (Wilcoxon p-value, 1.37 × 10^−6^). Importantly, this tubulointerstitial immune profile was also more significant when distinguishing nephrotic syndrome subjects – FSGS, MCD and membranous nephropathy (MN) – with complete remission from similar subjects without remission. Non-remitting samples showing higher expression of these immune genes (Wilcoxon p-value = 1.52 × 10^−4^) (**Fig. 4**).

**Fig. 4.**
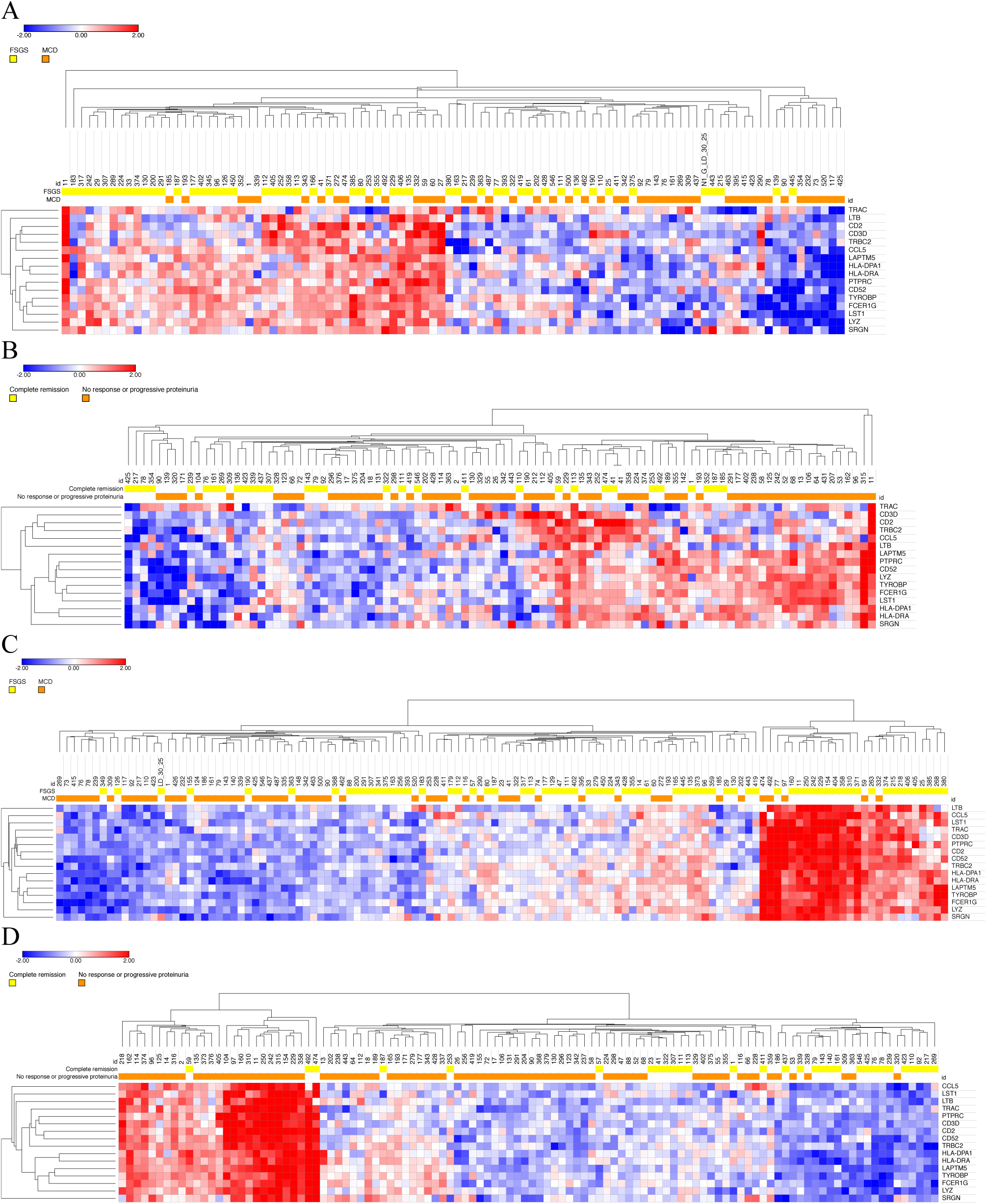
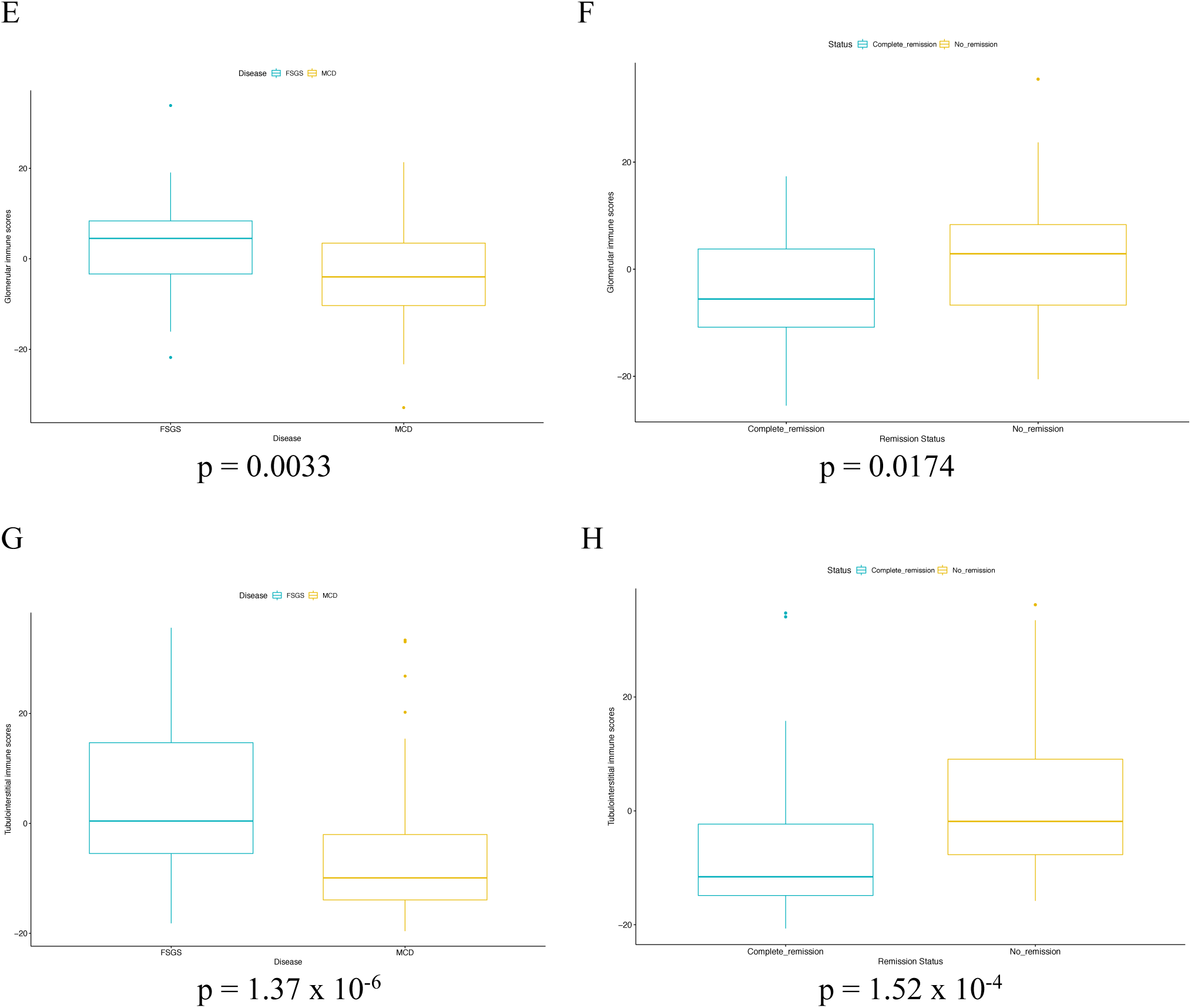
Heat maps and box plots showing the expression of the 16 most highly expressed genes from monocyte and lymphocyte clusters in the transcriptomic data from NEPTUNE cohort. **A-D**. Heat maps showing (**A**) Glomerular expression of MCD (n = 47) and FSGS (n = 51) samples (**B**) Glomerular expression of nephrotic syndrome samples (FSGS, MCD and membranous nephropathy (MN)) with complete remission (n = 31) and samples without remission (no response or progressive proteinuria) (n = 65) (**C**) Tubulointerstitial expression of MCD (n = 55) and FSGS (n = 68) samples (**D**) Tubulointerstitial expression of nephrotic syndrome samples with complete remission (n = 30) and samples without remission (n = 80). **E-H**. Box plots showing combined z-scores of 16 genes in (**E**) Glomerular expression data of MCD and FSGS samples (**F**) Glomerular expression data of all nephrotic syndrome samples with complete remission and samples without remission (**G**) Tubulointerstitial expression data of MCD and FSGS samples (**H**) Tubulointerstitial expression of all nephrotic syndrome samples with complete remission and samples without remission. The p-values for z-score comparisons were by Wilcoxon tests.

We evaluated whether the EMT signature was different between FSGS and MCD samples. The glomerular expression profile of the ten most highly expressed EMT-related genes in podocytes (**Table S13**) showed significant difference between the two diseases, with FSGS samples showing higher EMT signature. The Wilcoxon tests comparing the combined EMT signature scores were significant in both the glomerular compartment (Wilcoxon p-value = 2.0 × 10^−4^) and the tubulointerstitial compartment (Wilcoxon p-value = 3.35 × 10^−5^) (**Fig. 5**).

**Fig. 5.**
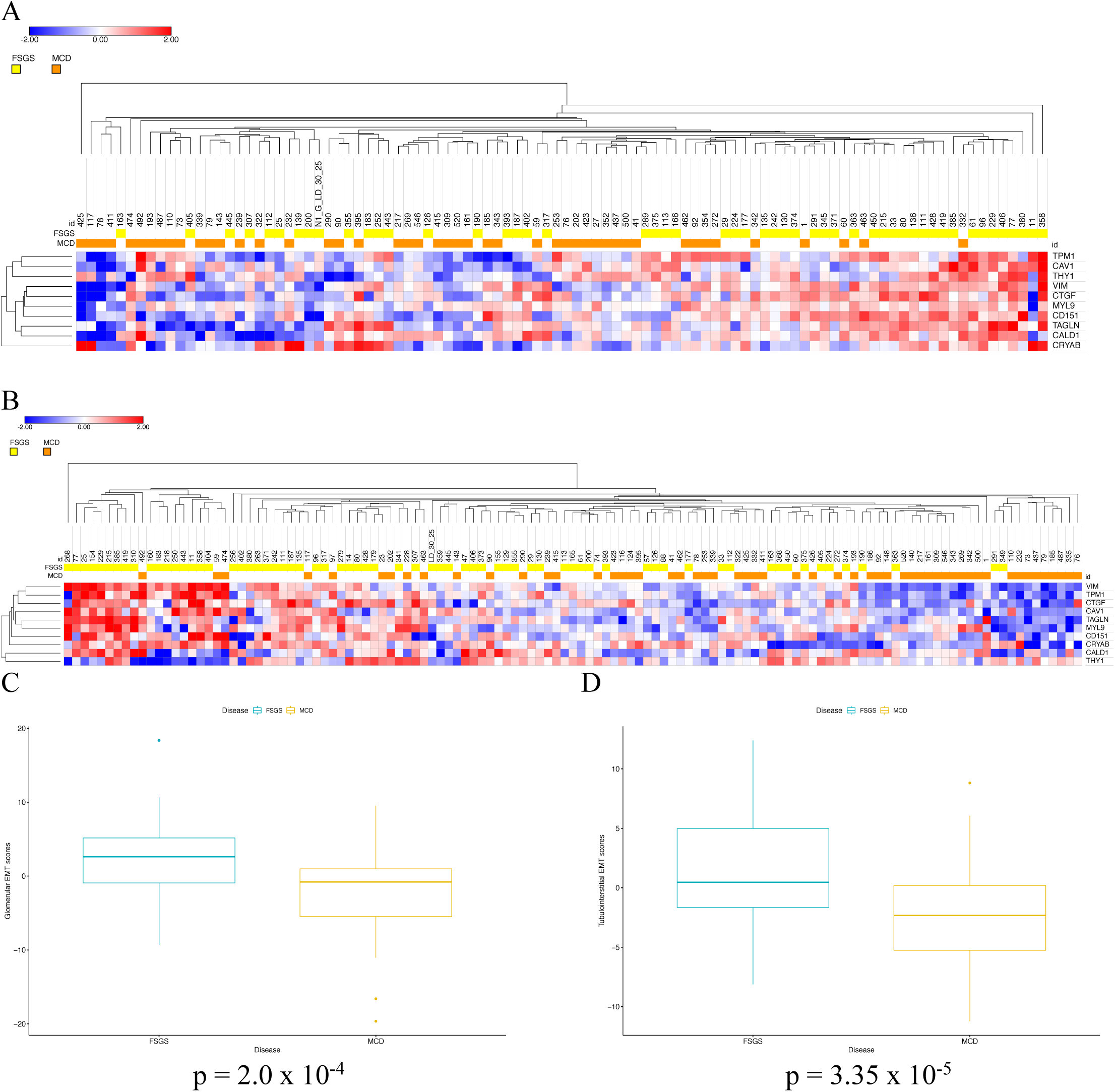
Heatmaps and boxplots showing the expression of the 10 EMT genes in the transcriptomic data from NEPTUNE cohort. (**A**) Heat map showing glomerular expression of MCD (n = 47) and FSGS (n = 51) samples (**B**) Heat map showing tubulointerstitial expression of MCD (n = 55) and FSGS (n = 68) samples (**C**) Box plot showing combined z-scores of 10 EMT genes in glomerular expression data of MCD and FSGS samples (**D**) Box plot showing combined z-scores of 10 EMT genes in tubulointerstitial expression data of MCD and FSGS samples.

### Immune signatures in lupus nephritis

To examine the universality of immune marker gene expression and their potential utility in other glomerular diseases, we evaluated their expression in the immune single cell data from the lupus nephritis subjects in the Accelerating Medicines Partnership (AMP) ^25^ as reported by Arazi et al ^26^. We observed that top 16 immune marker genes from FSGS monocytes and lymphocytes were also highly expressed in the monocyte and lymphocyte clusters, respectively from lupus nephritis subjects (**Fig. 6A and 6B**). We also evaluated the expression of these immune genes in the bulk RNA-seq data of urine samples from an American multi-ethnic cohort of lupus nephritis subjects and found that subjects with active disease (defined as urine protein: creatine ratio > 0.5) had higher expression of these genes than those with inactive disease (urine protein: creatine ratio ≤ 0.5), reflecting the higher presence of monocytes/lymphocytes in the urine of subjects with active lupus nephritis (**Fig. 6E**).

**Fig. 6.**
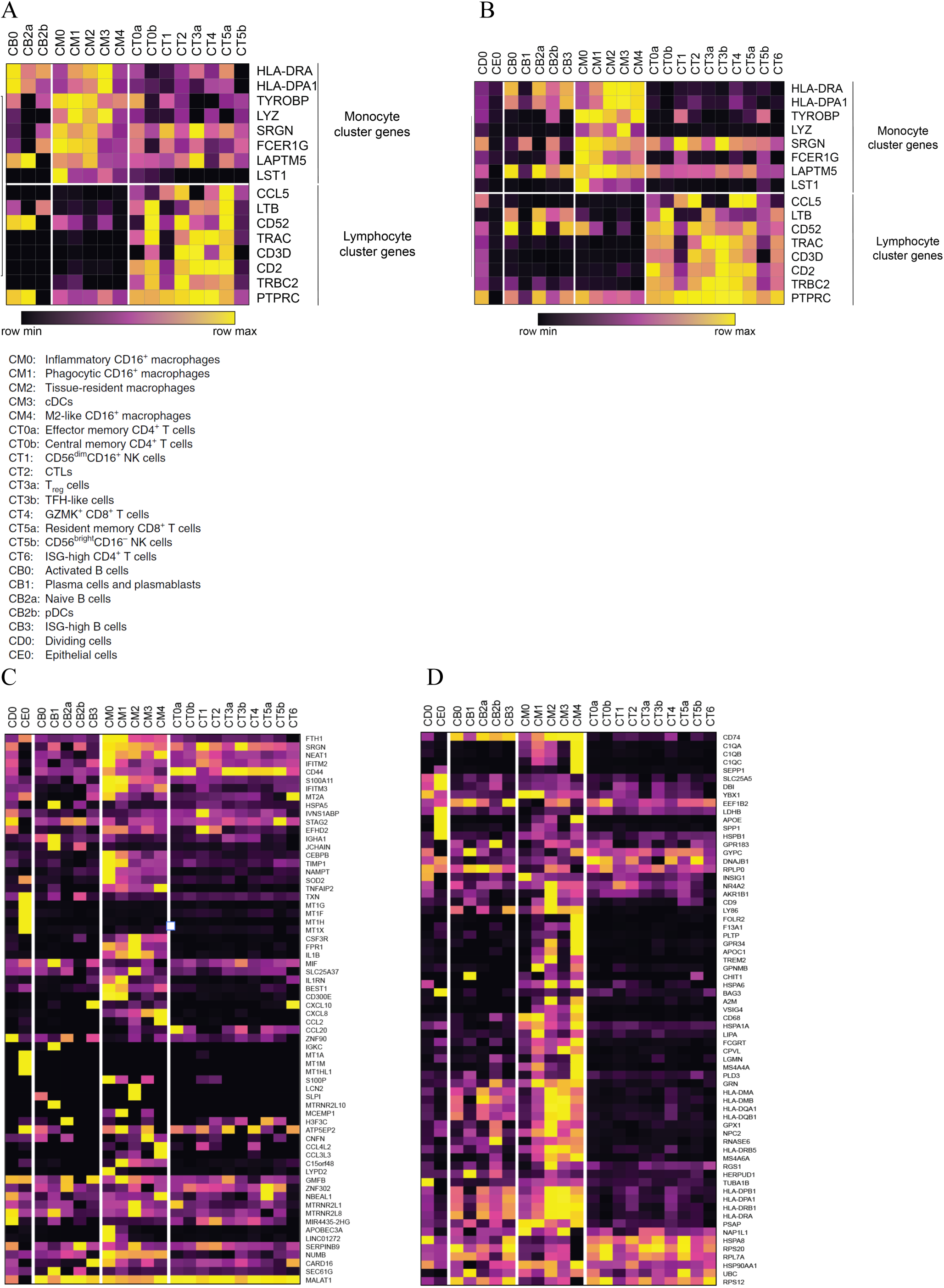
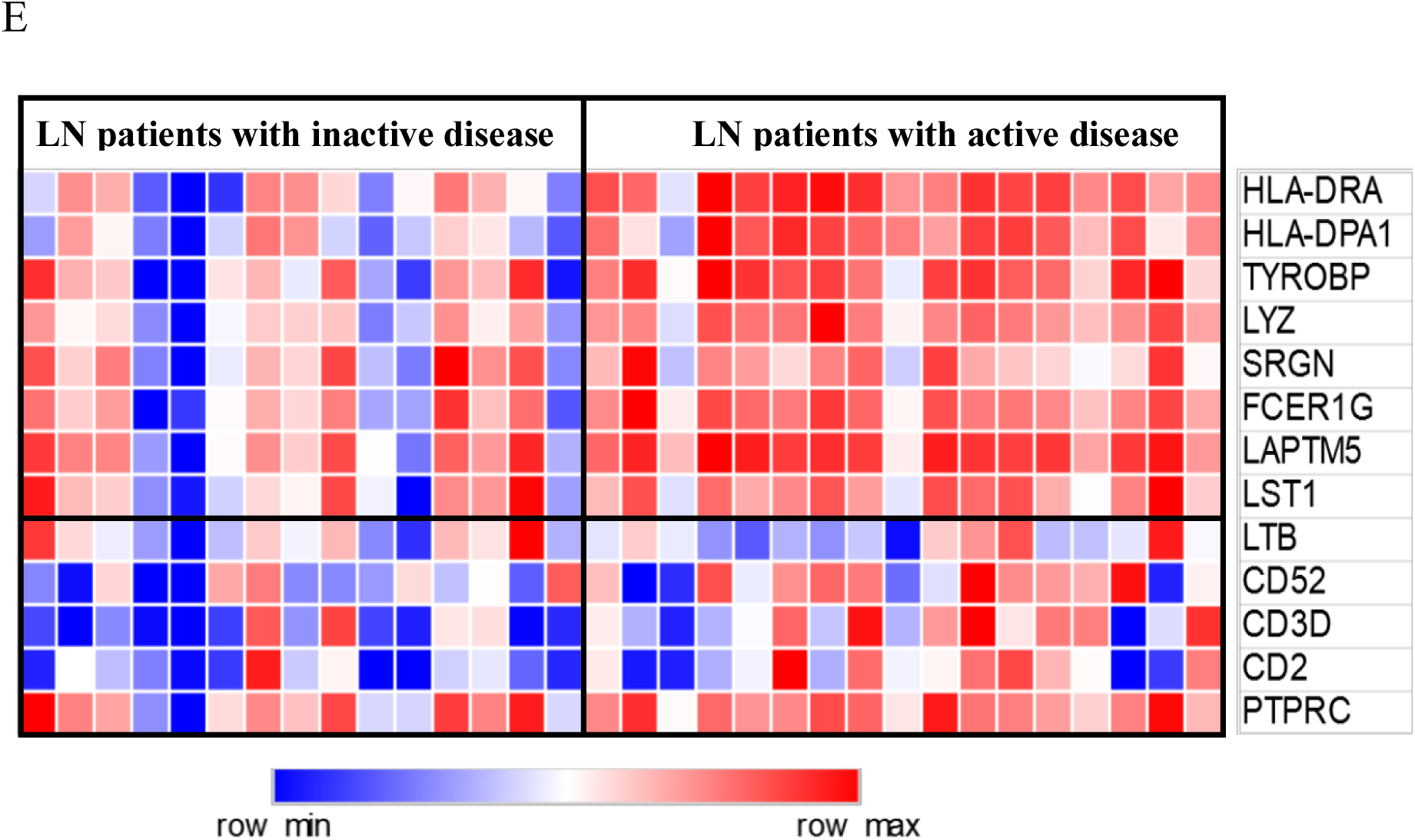
Expression of immune signature genes and M1 and M2 genes from FSGS urine single cell data in lupus nephritis. **A-B** Heatmaps showing the expression of the 16 most highly expressed genes from FSGS monocyte and lymphocyte clusters in the AMP single cell RNA-seq data of immune cells in lupus nephritis from (**A**) urine samples (**B**) kidney biopsy samples. **C-D** Heatmaps showing the expression of (**C**) M1 and (**D**) M2 genes from **Fig. 2C** in the AMP single cell RNA-seq data of immune cells from lupus nephritis kidney biopsy samples. (**E**) Heatmap showing the expression of the most highly expressed monocyte and lymphocyte markers in the urine bulk RNA-seq data of lupus nephritis patients with active and inactive disease.

Finally, we investigated the gene expression related to M1 and M2 signatures from the pseudotime heatmap (**Fig. 2C**) in the AMP kidney immune single cell data of lupus nephritis. The M1 signature genes were enriched in the myeloid cell clusters, especially in the inflammatory and phagocytic CD16^+^ macrophages (CM0 and CM1), and M2 signature genes were found to be more enriched in tissue resident macrophages, conventional dendritic cells and M2-like CD16^+^ macrophages (CM2, CM3 and CM4) (**Fig. 6C and 6D**).

## Discussion

In this study, we report on single-cell RNA sequencing results of urine cells from subjects with primary glomerular diseases, FSGS and MCD. This is, to our knowledge, the first attempt to comprehensively investigate the different cell types and their gene expression profiles in urine of such subjects. Our findings revealed a landscape of immune cells, podocytes, myofibroblasts and tubular cells with distinct expression profiles. We used canonical marker genes to identify the major cell types and confirmed those by annotation using Encode and Blueprint transcriptional reference data.

Urine podocytes showed loss of canonical podocyte markers such as *NPHS1, NPHS2* and *PODXL*, and high expression of markers for EMT (including mesenchyme and muscle markers). The protein products of these canonical genes are essential for the proper functioning of podocytes and their downregulation or loss may well contribute to podocyte injury, but that will require experimental confirmation. EMT is an important process in cancer biology, in which it is characterized by increased mobility of cancer cells. Podocytes may undergo a form of EMT ^27, 28^, leading to loss of differentiated function and possibly loss of the cells into the urinary space. The podocyte cluster was also positive for PEC markers, such as, *PAX2, PAX8* and *CLDN1* (**Fig. S3**) and it is possible to assume that there were some PECs in the podocyte cluster. However, PECs are also known to undergo EMT and these markers are also reported to be involved in the EMT process ^29, 30, 31, 32, 33^ and are also highly expressed in tubular and collecting ductal cell clusters, making it challenging to confirm their presence in the current study.

The presence of myofibroblasts in urine suggests that kidney cells are undergoing EMT and the resulting myofibroblasts may contribute to glomerulosclerosis and tubulointerstitial fibrosis. This is in part supported by annotation using Encode and Blueprint reference transcriptomic data, in which the podocyte and tubular cell clusters showed transcriptional similarities with myocytes and fibroblasts (**Fig. 2F**) and the higher EMT signature scores in FSGS samples from the NEPTUNE cohort. Tubular epithelial cell EMT has been proposed to be important for tubulointerstitial fibrosis in chronic kidney disease ^34, 35, 36^. In this study, we found transcriptional evidence of podocytes undergoing EMT in FSGS subjects.

Our analysis showed a variety of immune cells to be present in urine, predominantly monocytes, and we identified monocyte subtypes and characterized their gene expression profiles. Monocytes and/or macrophages expressing *APOE, APOC1* and *SPP1* have been reported in single cell studies of Alzheimer disease ^37^, atherosclerosis ^38^, and breast cancer ^14^. In atherosclerosis, these cells were considered to be foam cells due to their high expression of lipoproteins (*APOE* and *APOC1*). In our pseudotime analysis, we found that the FSGS monocytes constituted two branches, one with M1 characteristics (*TIMP1, IL1B* expression) and the other with M2 features (*APOE, APOC1* expression). Using the characteristic genes from each monocyte subset (*IL1B* for M1 and *APOE* for M2 monocytes), and examining publicly available single cell expression datasets, we observed a similar pattern of monocyte subtypes in kidney allograft rejection ^18^ and in cancers – melanoma ^15^, head and neck cancer ^16^. Additionally, the urinary M1- and M2-related signature genes from our pseudotime analysis were found to be enriched in the genes that are also expressed in myeloid sub-populations of kidney immune cells from lupus nephritis. This shared pattern across diverse diseases demonstrates that these inflammatory macrophage expression signatures are common across several inflammatory conditions with diverse etiologies. We also found a monocyte cluster with features of inflammatory dendritic cells. These cells are known to be involved in the initiation and maintenance of TH17 cell response, which have been implicated in several autoimmune and inflammatory diseases ^39, 40, 41^. The distinct sub-populations of myeloid cells in lupus nephritis suggests that the FSGS pseudotime analysis may have captured the general M1 and M2 polarization states and the analysis of leukocytes from FSGS kidney tissue samples may also reveal further sub-populations.

Expression of genes specifically enriched in urinary monocyte clusters could contribute to podocyte injury. This could be the downstream result of podocyte injury, or could be mechanistically independent of podocyte injury. In the case of the two former possibilities, they could serve as biomarkers to detect ongoing podocyte injury. We found that inflammatory monocytes expressed high levels of *PLAUR* (encoding suPAR), which is consistent with previous studies. These monocytes could, therefore, be a source of plasma and urinary suPAR, which has been implicated in FSGS pathogenesis. *APOE* was the most significantly upregulated gene in FSGS monocytes when compared with peripheral blood monocytes from a healthy individual. Serum and urine levels of *APOE* are elevated in FSGS and nephrotic syndrome ^42^.

Other top upregulated genes in FSGS monocytes included *SPP1* (encoding the immune modulator, osteopontin), *APOC1* and several metallothionein genes (*MT1G, MT2A, MT1X, MT1H, MT1E*, among others). Osteopontin is upregulated in several autoimmune and inflammatory diseases, including rheumatoid arthritis, multiple sclerosis, Crohn disease, cancers and atherosclerosis; targeting osteopontin by monoclonal antibodies in rheumatoid arthritis primate models ameliorated the symptoms ^43^.

Based on the expression profiles of the most highly expressed immune and EMT genes in FSGS urine single cell samples, we next evaluated renal expression of these genes in transcriptomic data from the NEPTUNE cohort of primary nephrotic disease. This analysis was informative for three reasons. First, NEPTUNE provided validation of our findings in a larger cohort of different sample type (kidney biopsy) and technology (microarray data). Second, the NEPTUNE cohort contains expression data from MCD biopsies, which is a major entity in the differential diagnosis of FSGS; this served as a disease control for FSGS. Third, it enabled us to correlate our gene expression data with nephrotic disease remission in this cohort.

The tubulointerstitial expression of the top monocyte/lymphocyte genes was higher in FSGS than in MCD samples and was also higher in nephrotic syndrome samples without remission than those in complete remission. This suggests that the active tubulointerstitial inflammation is important for the development of FSGS and the resistance to treatment.

Similarly, the expression of top immune signature genes was higher in the urine of subjects with active disease than those with inactive disease. This reflects the presence of higher number of immune cells in subjects with active nephritis and the potential for the use of these marker genes to monitor disease activity.

In the NEPTUNE data, both glomerular and tubulointerstitial expression of EMT genes were also significantly higher in FSGS than in MCD samples, likely reflecting the cell injury and fibrotic changes in FSGS which are not typically seen in MCD. Since the NEPTUNE EMT signature is from bulk expression data, the glomerular EMT signature can also be contributed by the parietal epithelial cells, which can also undergo EMT as in the setting of glomerulonephritis ^44, 45, 46^.

EMT is known to be induced by M2 monocytes through TGF-β signaling ^47, 48^. Our cell-to-cell interaction analysis showed *TGFB1* signaling between kidney epithelial cells and all immune cell types. It also revealed signaling of cytokines from TNF family. Recently, a study using the transcriptomic data from NEPTUNE and the European Renal cDNA Bank (ERCB) also identified high TNF activation signatures in a subset of nephrotic syndrome samples with mainly (∼80%) FSGS cases. The same group of subjects showed a higher risk of disease progression compared to other groups with low TNF signatures ^49^. Previously, however, treatment of therapy-resistant FSGS patients with adalimumab in the FONT trial (novel therapies in resistant FSGS) was proven unsuccessful, with only 2 patients showing dramatic improvement in proteinuria (from 17 to 0.6mg/mg and from 3.6 to 0.6mg/mg) ^49, 50^. The cell-cell interaction results from the current study also revealed that *TNFSF12*-*TNFRSF12A* (TWEAK/Fn14) and *TNFSF10*-*TNFRSF10B* (TRAIL/DR5) interactions are stronger than *TNF* interactions. Both TWEAK and TRAIL are known to induce apoptosis and were implicated in chronic inflammation, organ remodeling and fibrosis, and can be potential targets for immunotherapy ^51, 52, 53, 54, 55, 56, 57, 58^.

This study suggests that urine single-cell analysis might aid in the diagnosis and monitoring of kidney diseases. Here, scRNA-seq captured the expression landscape of renal epithelial cells undergoing EMT in the urine, which reflects the renal pathology of at least some FSGS patients. This approach also defined the gene expression profiles of immune cells in urine, identifying lymphocytes (which were mostly T cells) and macrophages (distinguishing between M1 and M2 monocytes). This approach could not, however, distinguish which compartment (glomerular or tubulointerstitial) the immune cells originated from and could not distinguish between podocytes from the sclerosed and non-sclerosed glomeruli of the same patient. These issues could be addressed by single cell spatial transcriptomic approaches using kidney biopsy samples.

Due to the limited number of urine single cell samples, we could not evaluate the diagnostic and prognostic potential of urine immune cells for FSGS. Studies with larger sample sizes will be needed to capture different immune cells in the urine, possibly by florescence-activated cell sorting (FACS) and determining their sensitivity and specificity for distinguishing FSGS from MCD or predicting FSGS remission. Similarly, the protein products of some of the top inflammatory genes from monocytes could be quantified by enzyme-linked immunosorbent assay (ELISA) in serum and urine of patients and evaluated for diagnostic and prognostic potential.

In summary, this study describes in detail the transcriptional profile of different cell types present in the urine of FSGS subjects and provides insights into relevant pathophysiological processes. We validated these findings in NEPTUNE renal transcriptomic data, where we found upregulation of selected immune and EMT genes in FSGS similar to what we had observed in urine immune cells and urine kidney epithelial cells, respectively. These findings suggest the possibility of using urine as the liquid biopsy, which, in contrast to kidney biopsy, could be repeated as needed. Finally, the data in this study suggests an important role of immune cells in the pathogenesis of glomerular disease and proposes potential disease biomarkers for further exploration.

## Methods

### Study design and sample preparation

We collected a total of 23 non-first-void morning urine samples from 12 FSGS patients. Out of 23 samples, seven samples were collected from one male African American subject over a period of six months, four samples from a female African American subject, two samples from another female African American subject and two samples from a male Asian subject (**Fig. S13** and **Table S14**). A clinical research protocol was approved in advance and all subjects provided written informed consent or assent.

### Urine sample processing, FACS sorting, and single cell capture

Subjects collected urine as a second void two to four hours after first voiding in the morning applying clean-catch practices. Whole urine (50 to 100 ml) samples were filtered (70 µm) and then sedimented for 10 to 15 min at 300 x g at 4 °C. The sediment was washed twice with ice-cold 0.04% bovine serum albumin in Dulbecco’s phosphate buffered saline and then either subjected to FACS sorting or to immediate cell capture using the 10x Genomics platform (**Fig. S14**). For FACS sorting, urine cells were buffer exchanged into Flow Cytometry Staining Buffer (eBioscience, Invitrogen, Carlsbad, CA), filtered through a 40 µm filter, combined with 7AAD and HOECHST and sorted for debris-free, nucleated live cells on a BDFASC Fusion (BD Biosciences, San Jose, CA) with a 70 µm nozzle size at 70 psi sheath pressure. HOECHST-7AAD-positive cells were sorted into DPBS with 10% FBS and then subjected to immediate cell capture. Urine cells were captured using the scRNA-seq using Chromium Single Cell 3’ Library & Gel Bead Kit, v2 (10x Genomics, Pleasanton, CA) and processed following exactly the supplier provided protocol. The resulting mRNA libraries were sequenced on NextSeq 500 (Illumina, San Diego, CA) platform with typical setup as a 26 cycles + 57 cycles non-symmetric run. Demultiplexing was done allowing 1 mismatch per barcode. Sequencing data were analyzed with the Cellranger v2.2.0 software (10x Genomics) applying default parameters. Results from the analysis are shown in **Fig. S15** and **Table S15**.

### Data integration and batch correction

Single cell gene expression data from all 23 samples were merged into a single dataset using Seurat (version 2.3.4, https://satijalab.org/seurat/). Cells with fewer than 100 detected genes or more than 6000 detected genes, and cells with mitochondrial transcript representing >10% of all transcripts were removed. Gene expression data was log normalized and scaled by regressing out the number of unique molecular identifiers and the mitochondrial percentage of the cells. We batch-corrected the data by the sample identities using the Harmony R package (version 0.1.0, Broad Institute, Boston, MA). Clustering of cells was done on batch-corrected data using 15 principal components that were calculated based on 6639 highly variable gene transcripts. In addition, we processed the gene expression data separately using Scanpy (1.4.4., https://github.com/theislab/scanpy) and employed two separate batch-effect minimization methods using either packages bbknn (https://github.com/Teichlab/bbknn) or scVI (https://github.com/YosefLab/scVI). All approaches led to similar results based on clustering and key gene transcripts indicating an overall robustness of Seurat and Scanpy-based methods.

### Calculating corrected expression levels from Harmony batch correction

The singular value decomposition of a matrix *M* is given by

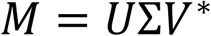

If n is the minimum and m the maximum of the number of rows and columns, U is the n by n matrix of embeddings, Σ is an n by m diagonal matrix of the singular values, and V is the m by m matrix of loadings. The embeddings in the Seurat object are actually *UΣ*.

The Harmony batch correction corrects the “PCs” (actually the cell embeddings). We followed the ansatz that Harmony is moving the cells around in PC space, without changing the SVD; i.e. as an approximation *Σ* and *U* are unchanged. Using this assumption, the original matrix *M* is approximated by

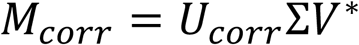

For the actual calculation we used a subset of the corrected PCs; an optimum reconstruction—as evaluated by the ability of the corrected gene expression levels to recapitulate the corrected PC after redoing the SVD—was obtained by using 900 PCs and 900 genes. The genes were the genes with the largest variation, augmented by the genes of particular interest for our analysis—i.e. genes identifying the cell types of interest. We used the R function propack.svd, in the R package svd (version 0.5) (https://CRAN.R-project.org/package=svd), for the SVD calculation; this approach requires more PCs than can readily be calculated by Seurat.

### scRNA-seq gene expression and pathway analysis

The most highly expressed genes in each cell cluster were identified by comparing gene expression of each cluster (podocytes, monocytes, lymphocytes, and tubular epithelial cells, and urothelial cells) with all the remaining cells in the dataset by Wilcoxon rank sum test. The three monocyte clusters were also combined to generate a list of expressed genes that could serve as marker genes for FSGS urine monocytes. Pathway analysis of monocyte and lymphocyte clusters was performed on the significantly expressed genes (adjusted p-value < 0.05) using Gene Ontology (GO) Biological Process library. For the podocyte cluster, due to the higher number of significantly expressed genes (n = 1492), we selected the top significant genes with log-fold change > 1 (n = 38 genes) and performed GO pathway analysis.

### Comparison and trajectory analysis of FSGS urine monocytes with peripheral blood monocytes from a healthy subject

The three monocyte clusters were extracted from the FSGS urine scRNA-seq data and merged with monocyte data from 10x Genomics (PBMC 8k v2 chemistry dataset) (https://support.10xgenomics.com/single-cell-gene-expression/datasets/2.1.0/pbmc8k?). The data integration and batch correction pipeline was applied again to the merged monocyte dataset. Genes in the FSGS monocytes whose expression was up-regulated compared to expression in PBMC monocytes were identified using the Wilcoxon rank sum test in Seurat.

The merged monocyte RNA expression dataset was imported into the Monocle2 R package (version 2.10.1, http://cole-trapnell-lab.github.io/monocle-release/) to carry out pseudotime trajectory analysis. Genes expressed in ≥5% of monocytes were selected for further analysis. We ran clustering and differential gene expression analysis across all clusters. We selected the 1500 most significant differentially expressed genes for dimension reduction and trajectory analysis. We ran the differentialGeneTest function in Monocle2 to identify genes that were differentially expressed along the trajectories as constructed by Mononcle2. We drew a heatmap We selected the most significant regulated genes with p-values less than 10^−20^. We assigned state 2 as the root state, since it contained all the peripheral blood monocytes from the healthy individual, which we considered to be not activated.

### Annotation of urine single cell clusters using Encode and Blueprint databases and the custom list of genes

We imported the whole urine cell Seurat dataset (or object) into the SingleR R package (version 1.0.1, https://bioconductor.org/packages/devel/bioc/html/SingleR.html). We annotated the FSGS urine single cell clusters using cell-type specific transcriptomic signatures from Encode and Blueprint databases. We also annotated the immune signatures of urine cells using the custom immune gene lists from Azizi et al. ^14^. Immune gene signatures were calculated for each cell and the signatures for each cluster was calculated as the mean of all the cells in that cluster. The R codes for these analyses were modified from the Github page (https://ncborcherding.github.io/files/CD8_Analysis.html).

### Evaluating expression of *IL1B, APOE* and *PLAUR* in the monocyte/macrophage data of publicly available scRNA-seq databases

We evaluated the genes that were most significantly differentially expressed between M1 and M2 monocytes in the publicly available single cell data. We selected *IL1B* as an M1 marker gene and *APOE* as an M2 marker gene and evaluated the expression of these two genes in macrophage data from melanoma (Jerby-Arnon et al, 2018) and from head and neck cancer (Puram et al, 2017), available at VirtualCytometry website (https://www.grnpedia.org/cytometry). The t-SNE plots of gene expression for these two selected genes were created on the interactive web page at that site and the differential expression tests of *APOE*^+^ macrophages compared with the all other macrophages were run using Wilcoxon rank sum test, in order to replicate M2 macrophage gene expression.

We evaluated the expression of *IL1B, APOE* and *PLAUR* in the kidney allograft rejection single cell data (Wu and Malone et al, 2018), available at Kidney Interactive Transcriptomics website (http://humphreyslab.com/SingleCell/). The t-SNE and violin plots were generated on the interactive web page.

### Cell-to-cell interaction analysis

We used CellPhoneDB ^22^ to investigate cell-to-cell interactions of ligand/receptor pairs among kidney cells and immune cells. The urogenital epithelial cell clusters were excluded from this analysis. Raw gene expression count data was taken from the Seurat object for kidney and immune cell clusters and normalized. Pairwise cluster-cluster interaction analyses were performed by randomly permuting the cluster labels for 10 times.

### Selecting the top immune and EMT genes and evaluating their expression in the NEPTUNE and lupus nephritis transcriptomic data

We selected the 16 most highly expressed genes in immune cells (8 genes from monocytes and 8 genes from lymphocytes) based on their high log-fold changes and the relatively low expression in the remaining cell clusters (**Table S13**). Similarly, we selected the top 10 most highly expressed EMT genes from the podocyte cluster (**Table S14**) and evaluated the expression of these genes in the kidney transcriptomic data from the NEPTUNE cohort. The NEPTUNE microarray transcriptomic data was log-normalized and batch-corrected using Combat. The values were converted to z-scores across all the samples in comparison and the z-scores of all the genes in each gene set were combined in each sample to compare between two groups by Wilcoxon rank sum tests. The heatmaps of gene expression were generated from z-scores using Morpheus https://software.broadinstitute.org/morpheus. We also evaluated the expression of the 16 genes in the urine and kidney immune single cell AMP Phase 1 lupus nephritis cohort (Arazi et al), as well as in the bulk RNAseq urine cells from an American multi-ethnic cohort (n= 17 active patients and 15 inactive patients) (unpublished). Heatmaps were generated using Morpheus from the Broad Institute (https://software.broadinstitute.org/morpheus/).

### Statistical Tests

Wilcoxon rank sum tests were used for the comparison of combined z-scores between NEPTUNE FSGS and MCD samples and the comparison between FSGS monocytes and healthy blood monocytes.

## Data Availability

Data is available from the authors on request during the review process and will be deposited to a public repository on publication.

## Acknowledgements

This work was supported by the Intramural Research Program of the NIDDK, NIH, Bethesda, MD (IRB protocols: 94-DK-0127 – Pathogenesis of Glomerulosclerosis; and 94-DK-0133 – FSGS Genetics). Support from CCR Single Cell Analysis Facility was funded by FNLCR Contract HHSN261200800001E, and support from Michigan Medicine was funded by O’Brien Kidney Research Core Center P30DK081943 (Matthias Kretzler). The Nephrotic Syndrome Study Network Consortium (NEPTUNE), U54-DK-083912, is a part of the National Institutes of Health (NIH) Rare Disease Clinical Research Network (RDCRN), supported through a collaboration between the Office of Rare Diseases Research, National Center for Advancing Translational Sciences and the National Institute of Diabetes, Digestive, and Kidney Diseases. Additional funding and/or programmatic support for this project has also been provided by the University of Michigan, the NephCure Kidney International and the Halpin Foundation. We thank Boehringer Ingelheim for supporting the establishment of the METAPHOR (MultiEThnic American luPus coHORt…) lupus nephritis cohort. We thank Bao Tran, Jyoti Shetty and other members of the CCR-Sequencing Facility for their help with the sequencing, Pradeep Kumar Dagur and Phil McCoy for their help with the FACS experiments, Luis Fernando Menezes for critical manuscript review and Jodi Blake, Noor Khalil and Tina Mainieri for helping with sample collection and data retrieval.

## Author contributions

J.B.K and C.A.W conceived the study. J.H designed the experiments and J.H, J.H.J and K.Z.L performed the experiments. M.C.K helped with single cell capture. M.M. and P.K. helped with the single-cell capture and next-generation sequencing. K.Z.L, J.H and J.H.J performed the data analysis. Y.Z, V.C, G.W.N and M.C gave advice on data analysis and G.W.N performed back-calculation of batch-corrected expression levels. C.C.B, S.E and M.K supported NEPTUNE transcriptomic data and C.C.B performed the analysis of lupus nephritis data. J.B.K, C.A.W, K.Z.L, J.H, A.Z.R, M.K and T.Y interpreted the results and K.Z.L prepared and wrote the manuscript.

## Materials and Correspondence

Khun Zaw Latt

## Supplementary Materials

### Members of the Nephrotic Syndrome Study Network (NEPTUNE)

**Fig. S1**. Stackplots showing the number of cells per cell type category.

**Fig. S2**. t-SNE plots showing expression of canonical marker genes for leukocytes, renal epithelial and urothelial cells.

**Fig. S3**. t-SNE plots showing expression of markers for PECs.

**Fig. S4**. Expression profiles of dendritic cell marker genes in the urine scRNA-seq data.

**Fig. S5**. PCs calculated from backcalculated expression levels preserve the Harmony batch correction.

**Fig. S6**. tSNE plots colored by back-calculated gene expression levels, for comparison with gene expression levels shown in **Fig. S1**.

**Fig. S7**. Trajectory plot of monocytes showing the original samples of the cells.

**Fig. S8**. The single cell RNA-seq data of macrophages from head and neck cancer (**A-C**) and melanoma (**D-F**) showing the expression of *IL1B* (representing M1) and *APOE* (representing M2).

**Fig. S9**. The single cell RNA-seq data from human kidney allograft rejection from Wu and Malone et al, retrieved from Humphreys Lab.

**Fig. S10**. High expression of *PLAUR* (encoding suPAR) found in monocyte clusters.

**Fig. S11**. Violin plots showing the most highly expressed monocyte/lymphocyte marker genes that were used to interrogate the presence of immune cells in the NEPTUNE transcriptomic data.

**Fig. S12**. Violin plots showing the most highly expressed monocyte/lymphocyte marker genes in the single nuclear RNAseq data from human adult kidney tissue from Menon et al.

**Fig. S13**. Stackplots showing cell population (%) by covariates.

**Fig. S14**. Overview of the experiments and analysis of the urine single cell study.

**Fig. S15**. Violin plots showing (**A**) number of genes (**B**) number of unique molecular identifiers and (**C**) mitochondrial percentage of individual clusters in the urine FSGS single cell dataset.

**Table S1**. The percentage of cells from each subject contributing to different cell clusters in the FSGS urine scRNA-seq study.

**Table S2**. The most highly expressed genes in podocyte cluster compared with remaining clusters.

**Table S3**. Gene ontology pathway analysis of the most highly expressed genes (log FC >= 1) from podocyte cluster (n = 38).

**Table S4**. The most highly expressed genes in monocyte clusters compared with remaining clusters.

**Table S5**. Gene ontology pathway analysis of significant genes from monocyte clusters (n = 817).

**Table S6**. The most highly expressed genes in lymphocyte cluster compared with remaining clusters.

**Table S7**. Gene ontology pathway analysis of significant genes from lymphocyte cluster (n = 481).

**Table S8**. Top 25 upregulated genes in *APOE*^+^ macrophages when compared with *APOE*^-^ macrophages in untreated melanoma from Jerby-Arnon et al.

**Table S9**. Top 25 upregulated genes in *APOE*^+^ macrophages when compared with *APOE*^-^ macrophages in head and neck cancer from Puram et al.

**Table S10**. Top 25 upregulated genes in *IL1B*^+^ macrophages when compared with *IL1B*^-^ macrophages in untreated melanoma from Jerby-Arnon et al.

**Table S11**. Top 17 upregulated genes in *IL1B*^+^ macrophages when compared with *IL1B*^-^ macrophages in head and neck cancer from Puram et al.

**Table S12**. The 16 most highly expressed genes from monocyte and lymphocyte clusters selected to be evaluated in the NEPTUNE kidney transcriptomic data.

**Table S13**. Ten genes from the podocyte cluster reported to be associated with EMT.

**Table S14**. Demographic characteristics of participants in the FSGS urine single cell RNA-seq study.

**Table S15**. Statistics of the cell counts, barcodes, reads, genes and UMIs for each sample.

## References

1. Mak SK, Short CD, Mallick NP. Long-term outcome of adult-onset minimal-change nephropathy. Nephrol Dial Transplant 11, 2192–2201 (1996).

2. Waldman M, et al. Adult minimal-change disease: clinical characteristics, treatment, and outcomes. Clin J Am Soc Nephrol 2, 445–453 (2007).

3. Park J, Liu C, Kim J, Susztak K. Understanding the kidney one cell at a time. Kidney International 96, 862–870 (2019).

4. Park J, et al. Single-cell transcriptomics of the mouse kidney reveals potential cellular targets of kidney disease. Science 360, 758–763 (2018).

5. Wilson PC, Humphreys BD. Kidney and organoid single-cell transcriptomics: the end of the beginning. Pediatr Nephrol, (2019).

6. Wu H, Humphreys BD. The promise of single-cell RNA sequencing for kidney disease investigation. Kidney Int 92, 1334–1342 (2017).

7. Butler A, Hoffman P, Smibert P, Papalexi E, Satija R. Integrating single-cell transcriptomic data across different conditions, technologies, and species. Nat Biotechnol 36, 411–420 (2018).

8. Korsunsky I, et al. Fast, sensitive and accurate integration of single-cell data with Harmony. Nat Methods, (2019).

9. Matsumoto T, et al. Proteomic analysis identifies insulin-like growth factor-binding protein-related protein-1 as a podocyte product. Am J Physiol Renal Physiol 299, F776– 784 (2010).

10. Zhang J, Liu J, Wu J, Li W, Chen Z, Yang L. Progression of the role of CRYAB in signaling pathways and cancers. Onco Targets Ther 12, 4129–4139 (2019).

11. Faure-Andre G, et al. Regulation of dendritic cell migration by CD74, the MHC class II-associated invariant chain. Science 322, 1705–1710 (2008).

12. Qiu X, et al. Reversed graph embedding resolves complex single-cell trajectories. Nat Methods 14, 979–982 (2017).

13. Aran D, et al. Reference-based analysis of lung single-cell sequencing reveals a transitional profibrotic macrophage. Nat Immunol 20, 163–172 (2019).

14. Azizi E, et al. Single-Cell Map of Diverse Immune Phenotypes in the Breast Tumor Microenvironment. Cell 174, 1293–1308 e1236 (2018).

15. Jerby-Arnon L, et al. A Cancer Cell Program Promotes T Cell Exclusion and Resistance to Checkpoint Blockade. Cell 175, 984–997 e924 (2018).

16. Puram SV, et al. Single-Cell Transcriptomic Analysis of Primary and Metastatic Tumor Ecosystems in Head and Neck Cancer. Cell 171, 1611–1624 e1624 (2017).

17. Kim K, Yang S, Ha SJ, Lee I. VirtualCytometry: a webserver for evaluating immune cell differentiation using single-cell RNA sequencing data. Bioinformatics 36, 546–551 (2020).

18. Wu H, et al. Single-Cell Transcriptomics of a Human Kidney Allograft Biopsy Specimen Defines a Diverse Inflammatory Response. Journal of the American Society of Nephrology 29, 2069–2080 (2018).

19. Wei C, et al. Circulating urokinase receptor as a cause of focal segmental glomerulosclerosis. Nat Med 17, 952–960 (2011).

20. Wu H, Uchimura K, Donnelly EL, Kirita Y, Morris SA, Humphreys BD. Comparative Analysis and Refinement of Human PSC-Derived Kidney Organoid Differentiation with Single-Cell Transcriptomics. Cell Stem Cell 23, 869–881 e868 (2018).

21. Efremova M, Vento-Tormo M, Teichmann SA, Vento-Tormo R. (2019).

22. Vento-Tormo R, et al. Single-cell reconstruction of the early maternal-fetal interface in humans. Nature 563, 347–353 (2018).

23. Menon R, et al. Single cell transcriptomics identifies focal segmental glomerulosclerosis remission endothelial biomarker. JCI Insight, (2020).

24. Gadegbeku CA, et al. Design of the Nephrotic Syndrome Study Network (NEPTUNE) to evaluate primary glomerular nephropathy by a multidisciplinary approach. Kidney Int 83, 749–756 (2013).

25. Hoover P, et al. Accelerating Medicines Partnership: Organizational Structure and Preliminary Data From the Phase 1 Studies of Lupus Nephritis. Arthritis Care Res (Hoboken) 72, 233–242 (2020).

26. Arazi A, et al. The immune cell landscape in kidneys of patients with lupus nephritis. Nat Immunol 20, 902–914 (2019).

27. Li Y, Kang YS, Dai C, Kiss LP, Wen X, Liu Y. Epithelial-to-mesenchymal transition is a potential pathway leading to podocyte dysfunction and proteinuria. Am J Pathol 172, 299–308 (2008).

28. Yamaguchi Y, et al. Epithelial-mesenchymal transition as a potential explanation for podocyte depletion in diabetic nephropathy. Am J Kidney Dis 54, 653–664 (2009).

29. Di Palma T, Lucci V, de Cristofaro T, Filippone MG, Zannini M. A role for PAX8 in the tumorigenic phenotype of ovarian cancer cells. BMC Cancer 14, 292 (2014).

30. Li L, Wu Y, Yang Y. Paired box 2 induces epithelial-mesenchymal transition in normal renal tubular epithelial cells of rats. Mol Med Rep 7, 1549–1554 (2013).

31. Lv J, Sun B, Mai Z, Jiang M, Du J. CLDN-1 promoted the epithelial to migration and mesenchymal transition (EMT) in human bronchial epithelial cells via Notch pathway. Mol Cell Biochem 432, 91–98 (2017).

32. Ohse T, et al. The enigmatic parietal epithelial cell is finally getting noticed: a review. Kidney Int 76, 1225–1238 (2009).

33. Suh Y, et al. Claudin-1 induces epithelial-mesenchymal transition through activation of the c-Abl-ERK signaling pathway in human liver cells. Oncogene 32, 4873–4882 (2013).

34. Liu Y. Epithelial to mesenchymal transition in renal fibrogenesis: pathologic significance, molecular mechanism, and therapeutic intervention. J Am Soc Nephrol 15, 1–12 (2004).

35. Ng YY, et al. Tubular epithelial-myofibroblast transdifferentiation in progressive tubulointerstitial fibrosis in 5/6 nephrectomized rats. Kidney Int 54, 864–876 (1998).

36. Abbate M, Zoja C, Rottoli D, Corna D, Tomasoni S, Remuzzi G. Proximal tubular cells promote fibrogenesis by TGF-beta1-mediated induction of peritubular myofibroblasts. Kidney Int 61, 2066–2077 (2002).

37. Mathys H, et al. Single-cell transcriptomic analysis of Alzheimer’s disease. Nature 570, 332–337 (2019).

38. Fernandez DM, et al. Single-cell immune landscape of human atherosclerotic plaques. Nat Med 25, 1576–1588 (2019).

39. Hu Y, Shen F, Crellin NK, Ouyang W. The IL-17 pathway as a major therapeutic target in autoimmune diseases. Ann N Y Acad Sci 1217, 60–76 (2011).

40. Segura E, et al. Human inflammatory dendritic cells induce Th17 cell differentiation. Immunity 38, 336–348 (2013).

41. Segura E, Amigorena S. Identification of human inflammatory dendritic cells. Oncoimmunology 2, e23851 (2013).

42. Bruschi M, et al. Apolipoprotein E in idiopathic nephrotic syndrome and focal segmental glomerulosclerosis. Kidney Int 63, 686–695 (2003).

43. Lund SA, Giachelli CM, Scatena M. The role of osteopontin in inflammatory processes. J Cell Commun Signal 3, 311–322 (2009).

44. Fujigaki Y, et al. Mechanisms and kinetics of Bowman’s epithelial-myofibroblast transdifferentiation in the formation of glomerular crescents. Nephron 92, 203–212 (2002).

45. Ng YY, et al. Glomerular epithelial-myofibroblast transdifferentiation in the evolution of glomerular crescent formation. Nephrol Dial Transplant 14, 2860–2872 (1999).

46. Shimizu M, et al. Role of integrin-linked kinase in epithelial-mesenchymal transition in crescent formation of experimental glomerulonephritis. Nephrol Dial Transplant 21, 2380–2390 (2006).

47. Yao RR, Li JH, Zhang R, Chen RX, Wang YH. M2-polarized tumor-associated macrophages facilitated migration and epithelial-mesenchymal transition of HCC cells via the TLR4/STAT3 signaling pathway. World J Surg Oncol 16, 9 (2018).

48. Zhu L, Fu X, Chen X, Han X, Dong P. M2 macrophages induce EMT through the TGF-beta/Smad2 signaling pathway. Cell Biol Int 41, 960–968 (2017).

49. Mariani LH, et al., (2018).

50. Joy MS, et al. Phase 1 trial of adalimumab in Focal Segmental Glomerulosclerosis (FSGS): II. Report of the FONT (Novel Therapies for Resistant FSGS) study group. Am J Kidney Dis 55, 50–60 (2010).

51. Burkly LC. TWEAK/Fn14 axis: the current paradigm of tissue injury-inducible function in the midst of complexities. Semin Immunol 26, 229–236 (2014).

52. Burkly LC, Michaelson JS, Hahm K, Jakubowski A, Zheng TS. TWEAKing tissue remodeling by a multifunctional cytokine: role of TWEAK/Fn14 pathway in health and disease. Cytokine 40, 1–16 (2007).

53. Ucero AC, et al. TNF-related weak inducer of apoptosis (TWEAK) promotes kidney fibrosis and Ras-dependent proliferation of cultured renal fibroblast. Biochim Biophys Acta 1832, 1744–1755 (2013).

54. Wynn TA, Ramalingam TR. Mechanisms of fibrosis: therapeutic translation for fibrotic disease. Nat Med 18, 1028–1040 (2012).

55. Braithwaite AT, Marriott HM, Lawrie A. Divergent Roles for TRAIL in Lung Diseases. Front Med (Lausanne) 5, 212 (2018).

56. Haw TJ, et al. A pathogenic role for tumor necrosis factor-related apoptosis-inducing ligand in chronic obstructive pulmonary disease. Mucosal Immunol 9, 859–872 (2016).

57. Morissette MC, Parent J, Milot J. The emphysematous lung is abnormally sensitive to TRAIL-mediated apoptosis. Respir Res 12, 105 (2011).

58. Wu Y, et al. Increased serum TRAIL and DR5 levels correlated with lung function and inflammation in stable COPD patients. Int J Chron Obstruct Pulmon Dis 10, 2405–2412 (2015).

