## Supplementary material for "Urine single cell RNA-sequencing in focal segmental glomerulosclerosis reveals inflammatory signatures in immune cells and podocytes": Supplementary materials.pdf

**Members of the Nephrotic Syndrome Study Network (NEPTUNE)**

NEPTUNE Enrolling Centers

*Cleveland Clinic, Cleveland, OH*: K Dell*, J Sedor**, M Schachere#, J Negrey#
*Children's Hospital, Los Angeles, CA*: K Lemley*, E Lim#
*Children's Mercy Hospital, Kansas City, MO*: T Srivastava*, A Garrett#
*Cohen Children's Hospital, New Hyde Park, NY:* C Sethna*, K Laurent #
*Columbia University, New York, NY:* G Appel*, A Pradhan#
*Emory University, Atlanta, GA:* L Greenbaum*, C Wang**, C Kang#
*Harbor-University of California Los Angeles Medical Center:* S Adler*, J LaPage#
*John H. Stroger Jr. Hospital of Cook County, Chicago, IL:* A Athavale*, M Itteera
*Johns Hopkins Medicine, Baltimore, MD:* M Atkinson*, S Boynton#
*Mayo Clinic, Rochester, MN:* F Fervenza*, M Hogan**, J Lieske*, V Chernitskiy#
*Montefiore Medical Center, Bronx, NY:* F Kaskel*, M Ross*, P Flynn#
*NIDDK Intramural, Bethesda MD:* J Kopp*, J Blake#
*New York University Medical Center, New York, NY:* H Trachtman*, O Zhdanova**, F Modersitzki#, S Vento#
*Stanford University, Stanford, CA:* R Lafayette*, K Mehta#
*Temple University, Philadelphia, PA:* C Gadegbeku*, S Quinn-Boyle#
*University Health Network Toronto:* M Hladunewich**, H Reich**, P Ling#, M Romano#
*University of Miami, Miami, FL:* A Fornoni*, C Bidot#
*University of Michigan, Ann Arbor, MI:* M Kretzler*, D Gipson*, A Williams#, J LaVigne#
*University of North Carolina, Chapel Hill, NC:* V Derebail*, K Gibson*, E Cole#, J Ormond-Foster#
*University of Pennsylvania, Philadelphia, PA:* L Holzman*, K Meyers**, K Kallem#, A Swenson#
*University of Texas Southwestern, Dallas, TX:* K Sambandam*, Z Wang#, M Rogers#
*University of Washington, Seattle, WA:* A Jefferson*, S Hingorani**, K Tuttle**§, M Bray #, M Kelton#, A Cooper#§
*Wake Forest University Baptist Health, Winston-Salem, NC:* JJ Lin*, Stefanie Baker#

*Data Analysis and Coordinating Center*: M Kretzler, L Barisoni, J Bixler, H Desmond, S Eddy, C Gadegbeku, B Gillespie, D Gipson, L Holzman, V Kurtz, M Larkina, J Lavigne, S Li, CC Lienczewski, J Liu, T Mainieri, L Mariani, M Sampson, M Wladkowski, A Williams, J Zee

*Digital Pathology Committee*: Carmen Avila-Casado (UHN-Toronto), Serena Bagnasco (Johns Hopkins), Joseph Gaut (Washington U), Stephen Hewitt (National Cancer Institute), Jeff Hodgin (University of Michigan), Kevin Lemley (Children's Hospital LA), Laura Mariani (University of Michigan), Matthew Palmer (U Pennsylvania), Avi Rosenberg (NIDDK), Virginie Royal (Montreal), David Thomas (University of Miami), Jarcy Zee (Arbor Research) Co-Chairs: Laura Barisoni (Duke University) and Cynthia Nast (Cedar Sinai)

*Principal Investigator; **Co-investigator; #Study Coordinator
§Providence Medical Research Center, Spokane, WA

### Supplementary Figures

A

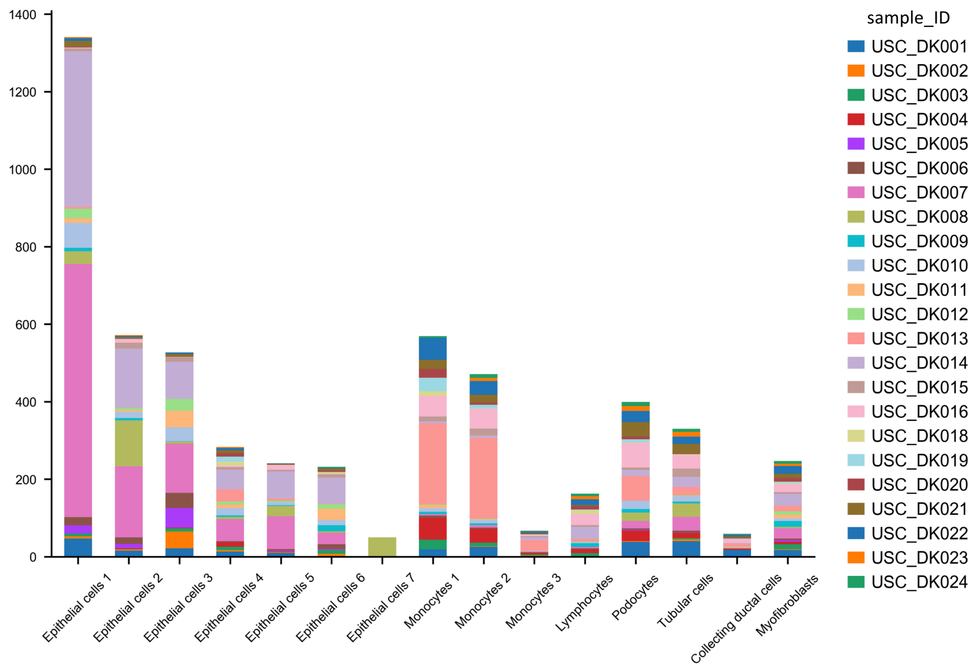


B

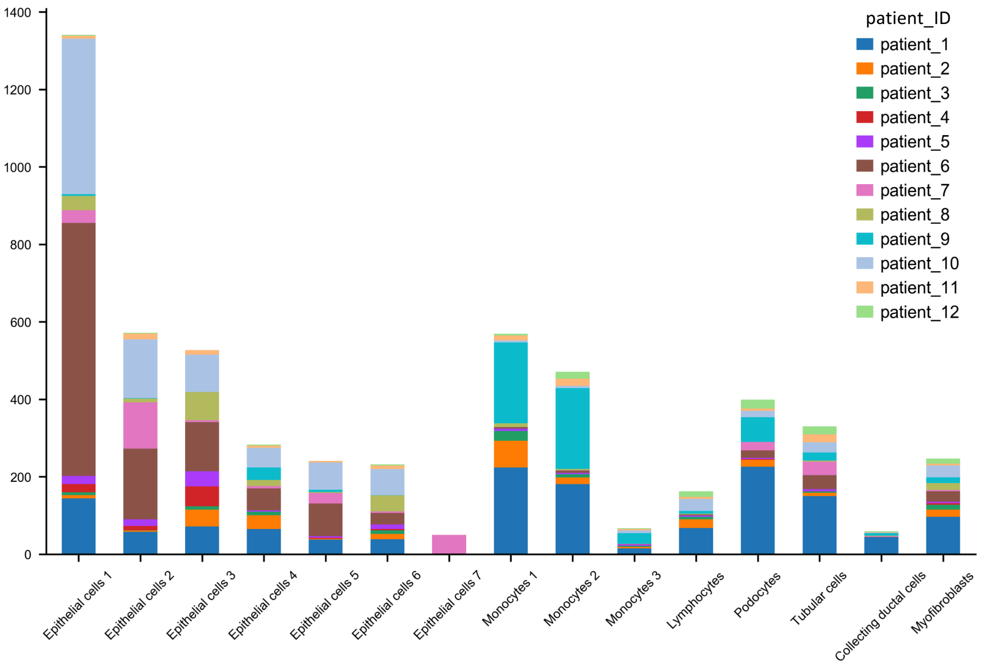


**Fig. S1.** Stackplots showing the number of cells per cell type category. (**A**) by sample ID (**B**) by patient ID.

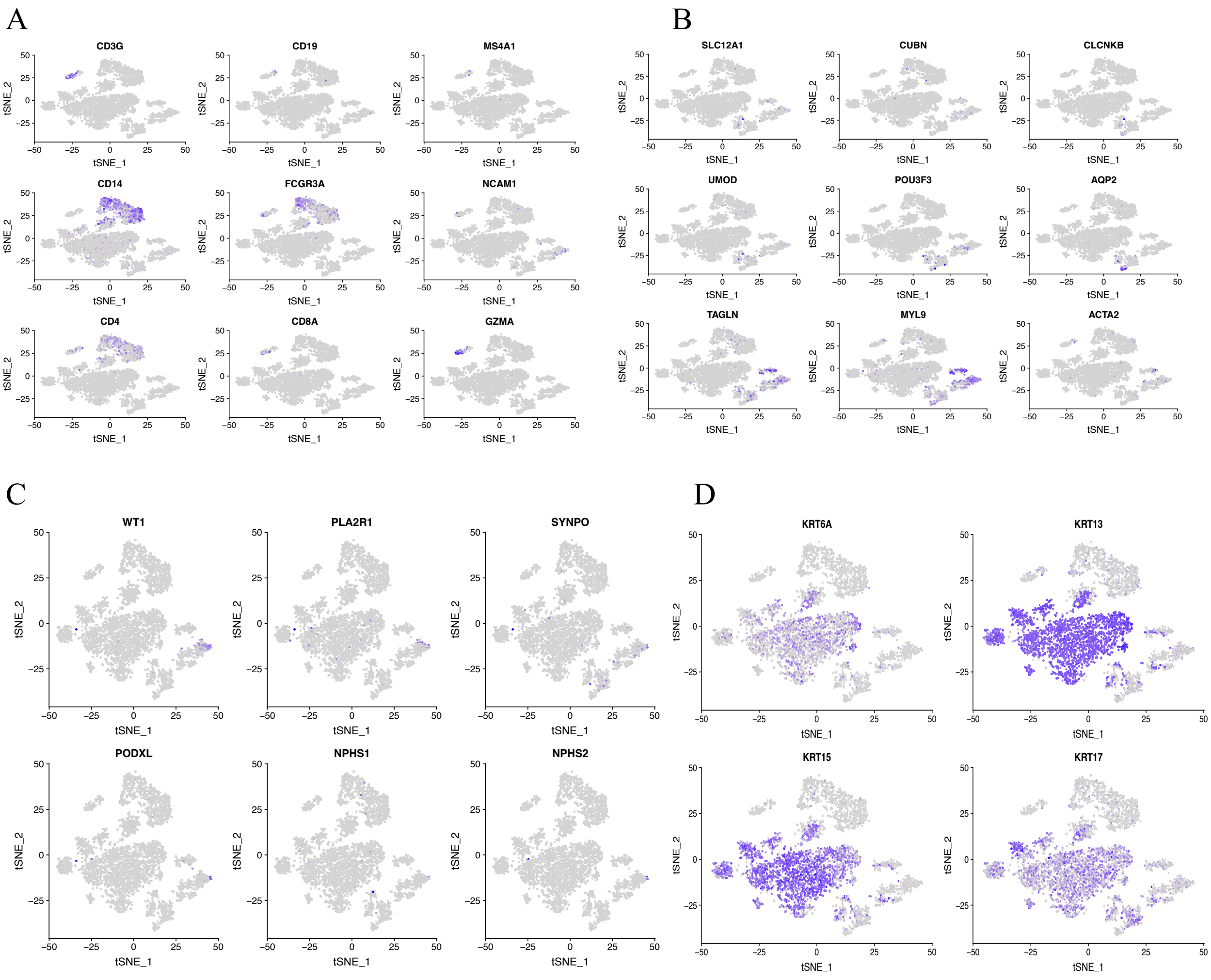


**Fig. S2.** t-SNE plots showing expression of canonical marker genes for leukocytes, renal epithelial and urothelial cells. (**A**) Shown are markers for T cells (*CD3G*, *CD4*, *CD8A* and *GZMA*), B cells (*CD19* and *MS4A1*), monocytes (*CD14* and *FCGR3A*) and NK cells (*NCAM1*). (**B**) Shown are markers for proximal tubular cells (*SLC12A1*, *CUBN*), distal tubular cells (*CLCNKB*), loop of Henle (*UMOD*, *POU3F3*), collecting duct (*AQP2*) and myofibroblasts (*TAGLN*, *MYL9* and *ACTA2*). (**C**) Shown are markers for podocytes and (**D**) markers for urogenital epithelial cells.

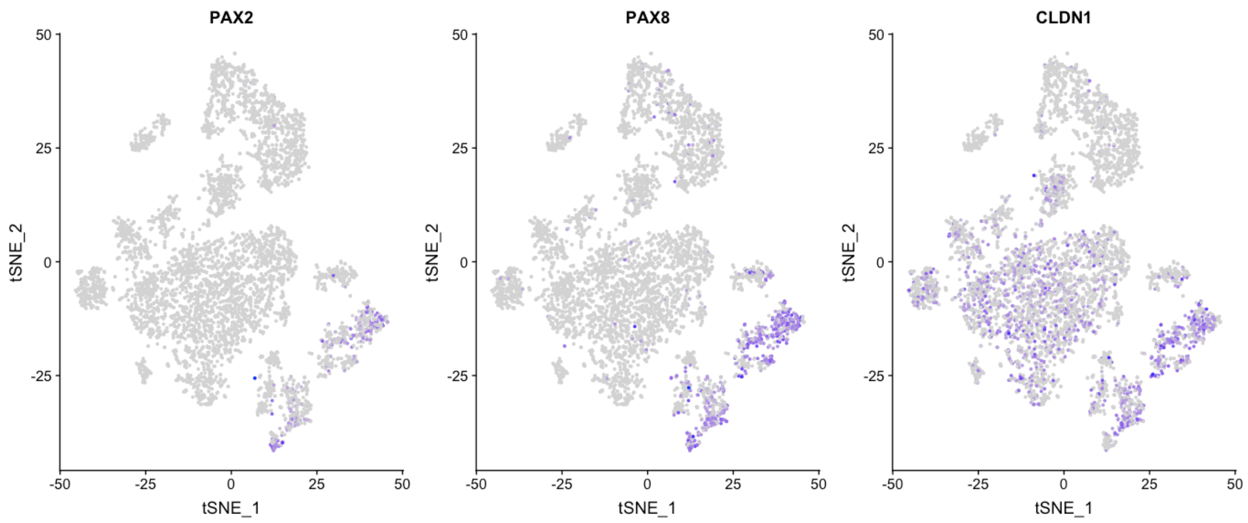


**Fig. S3.** t-SNE plots showing expression of markers for PECs.

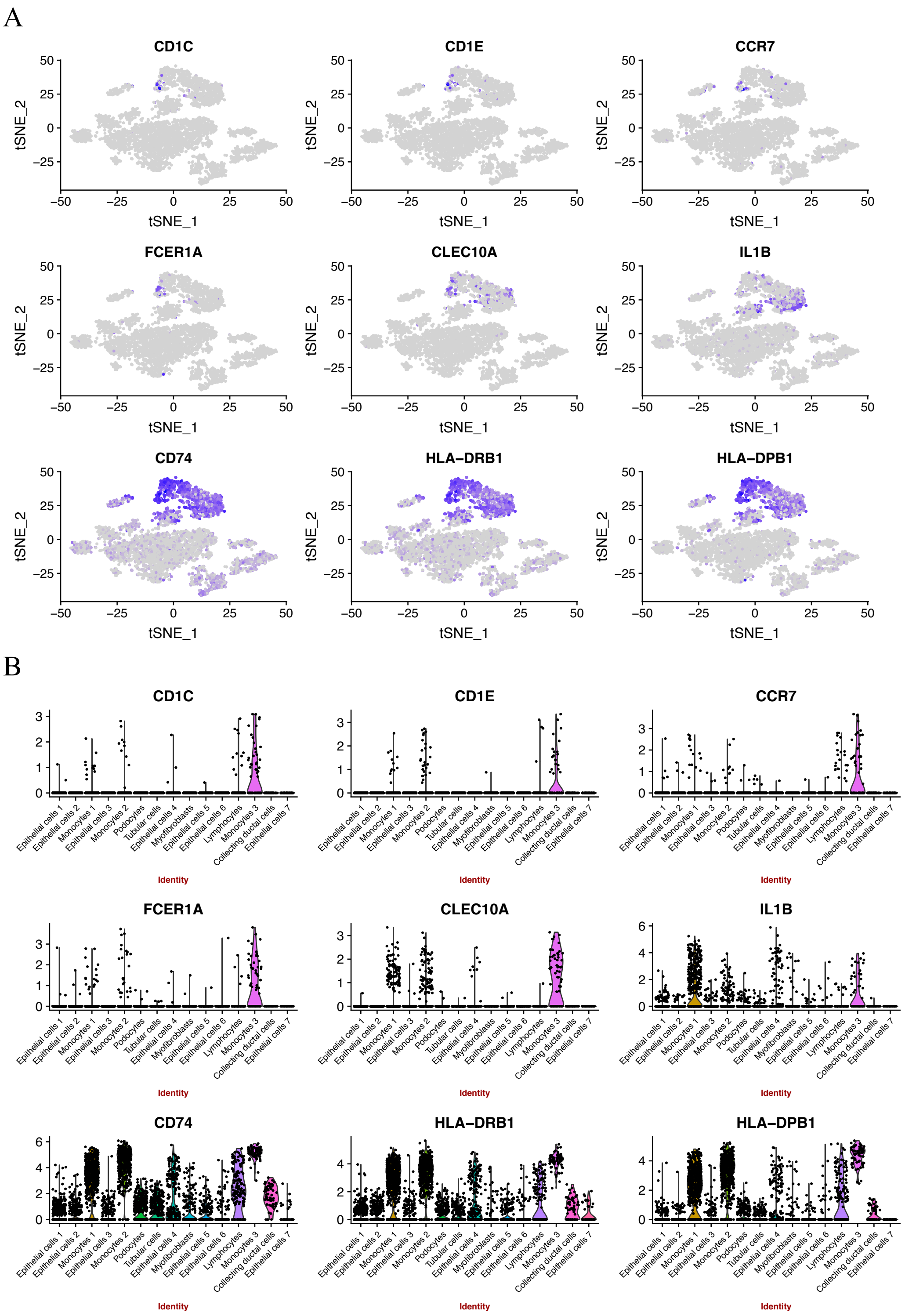


**Fig. S4.** Expression profiles of dendritic cell marker genes in the urine scRNA-seq data. (**A**) t-SNE plots and (**B**) violin plots. Cells in monocyte cluster 3 (Monocyte 3) had the highest expression levels of the dendritic cell marker genes.

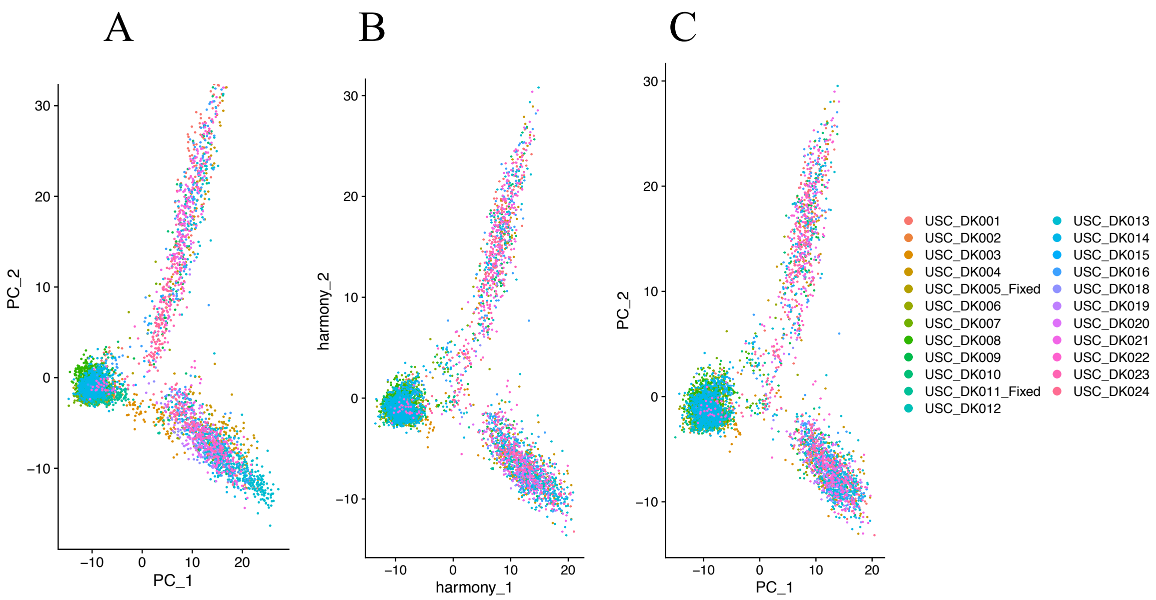


**Fig. S5.** PCs calculated from backcalculated expression levels preserve the Harmony batch correction. **(A)** Uncorrected PCs calculated by SVD from the Seurat SCT normalized expression levels, colored by batch (urine cell sample). The batch effect is shown most strongly by the blue tail on the lower right. **(B)** the same, corrected with Harmony. **(C)** Uncorrected PCs, from an SVD from SCT expression levels backcalculated from the Harmony corrected PCs (B); these are virtually identical to the Harmony corrected PCs. Thus the backcalculated SCT levels are effectively batch corrected.

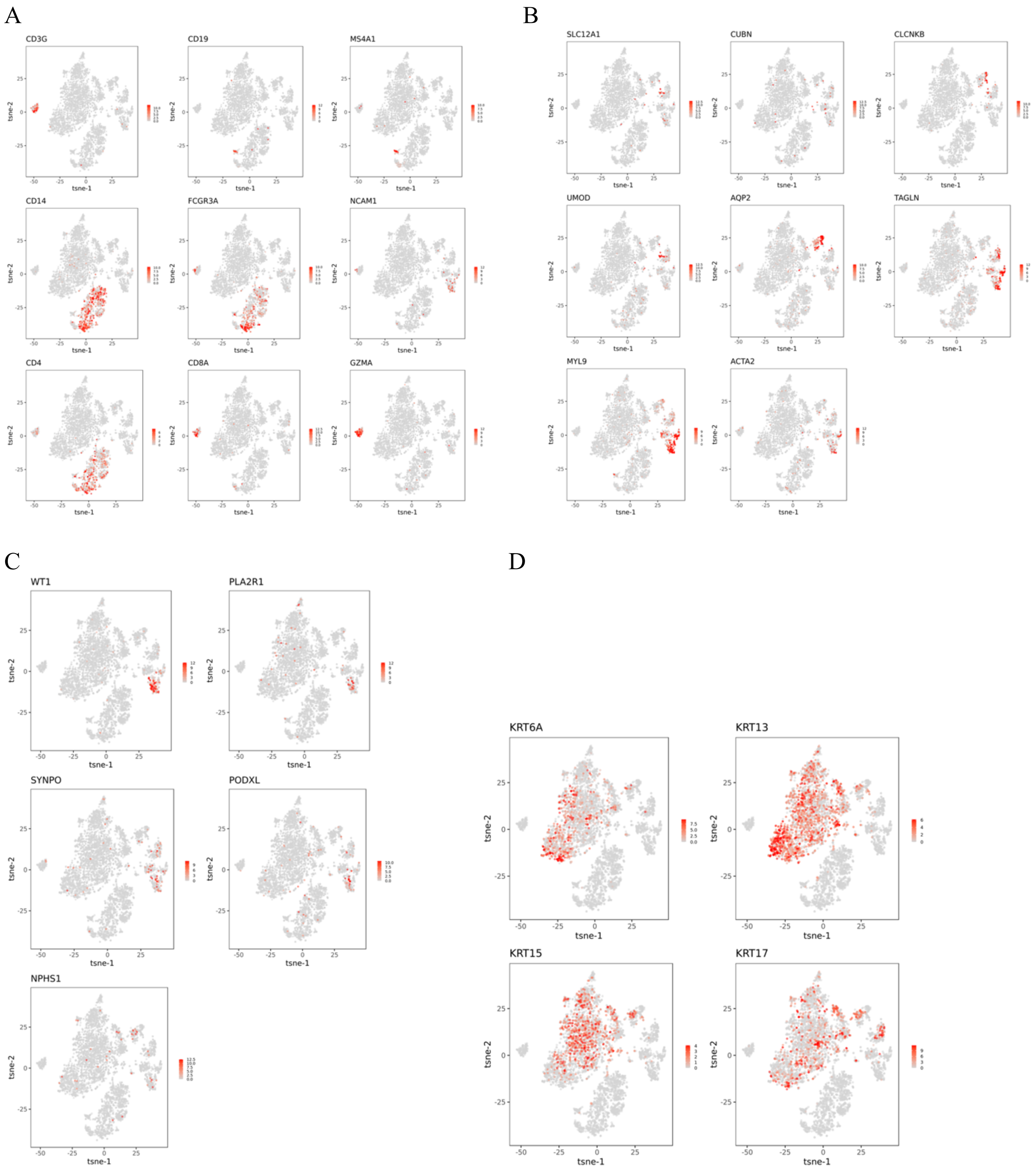


**Fig. S6.** tSNE plots colored by back-calculated gene expression levels, for comparison with gene expression levels shown in **Fig. S1**. **A, B, C, D:** back-calculated expression levels for genes shown in **Fig. S1A, B, C, D**.

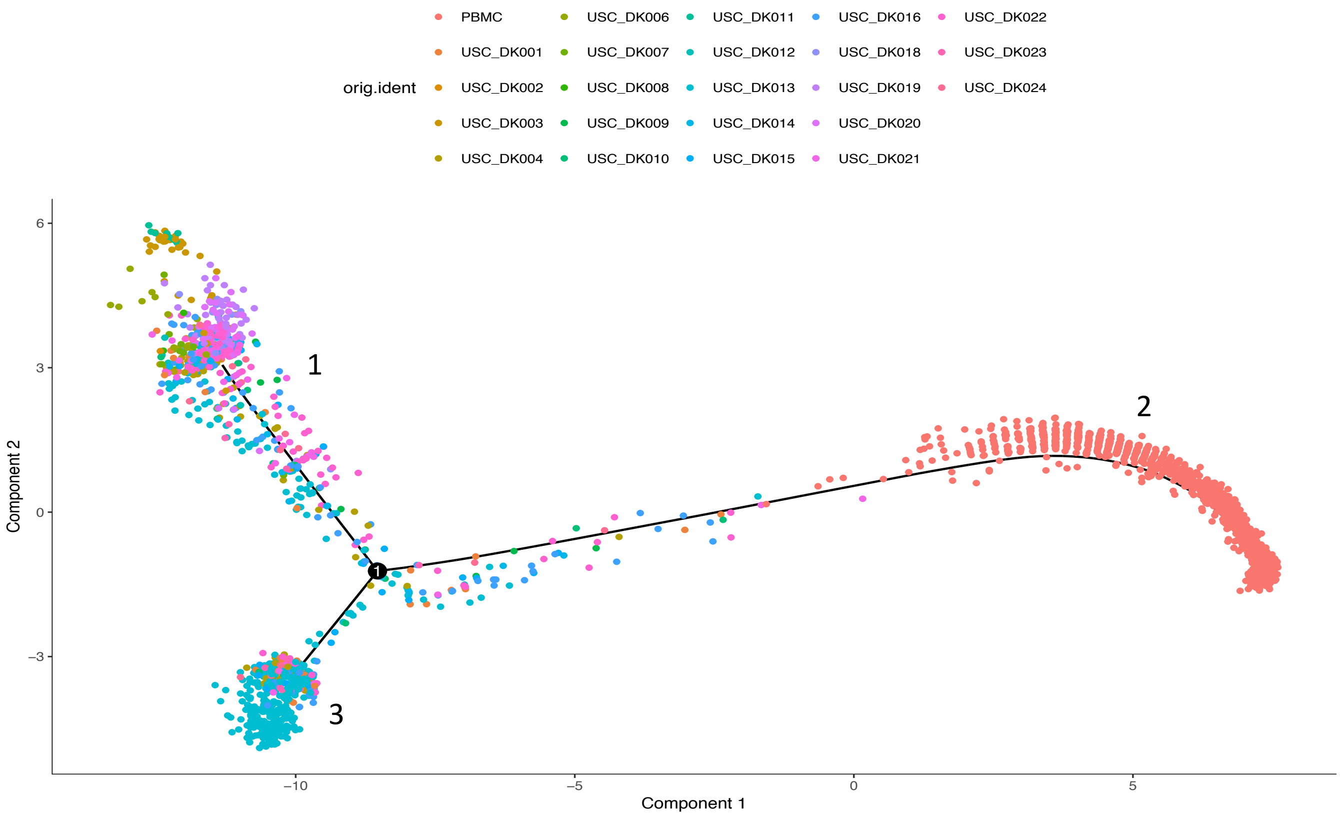


**Fig. S7.** Trajectory plot of monocytes showing the original samples of the cells. The peripheral blood monocytes from the 10x Genomics sample (PBMC) are concentrated in branch 2.

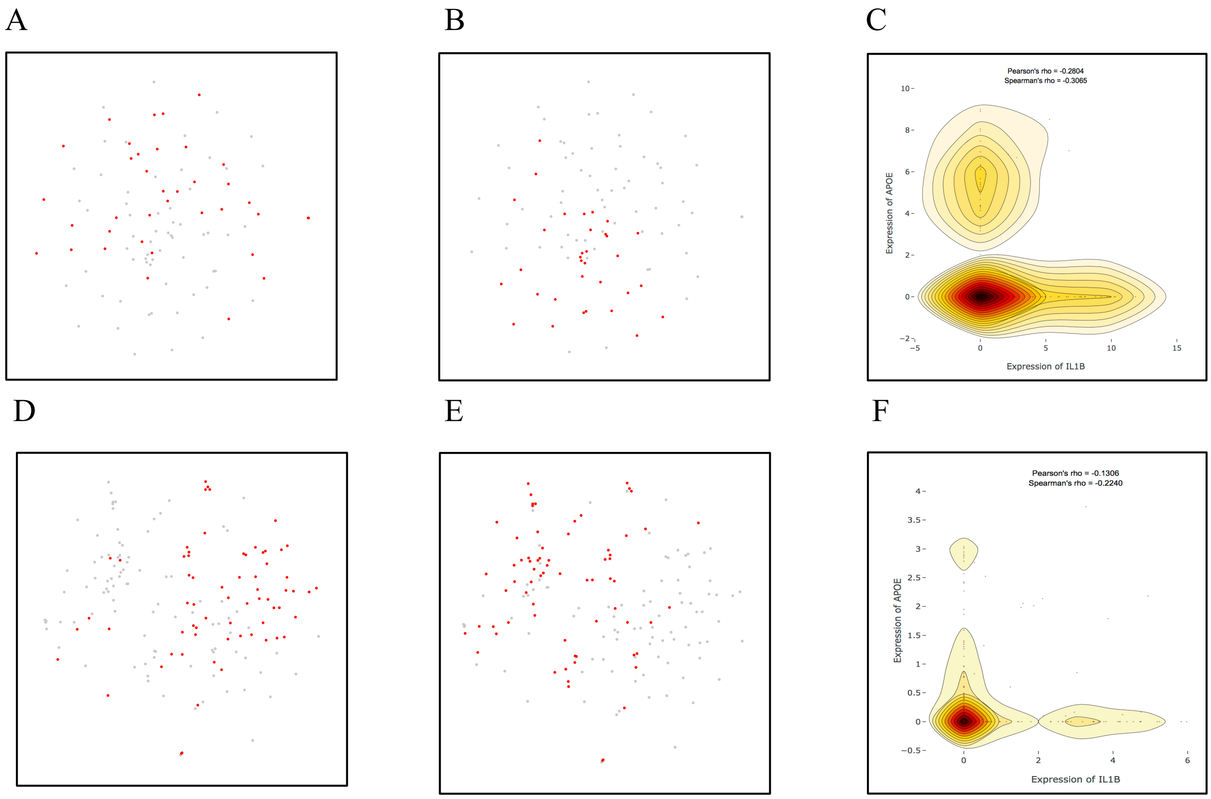


**Fig. S8.** The single cell RNA-seq data of macrophages from head and neck cancer (**A-C**) and melanoma (**D-F**) showing the expression of *IL1B* (representing M1) and *APOE* (representing M2). (**A**) t-SNE plot showing the expression of *IL1B* in head and neck cancer. (**B**) t-SNE plot showing the expression of *APOE* in head and neck cancer. (**C**) Correlation of the expression of *IL1B* and *APOE* in macrophages from head and neck cancer. (**D**) t-SNE plot showing the expression of *IL1B* in melanoma. (**E**) t-SNE plot showing the expression of *APOE* in melanoma. (**F**) Correlation of the expression of *IL1B* and *APOE* in macrophages from melanoma.

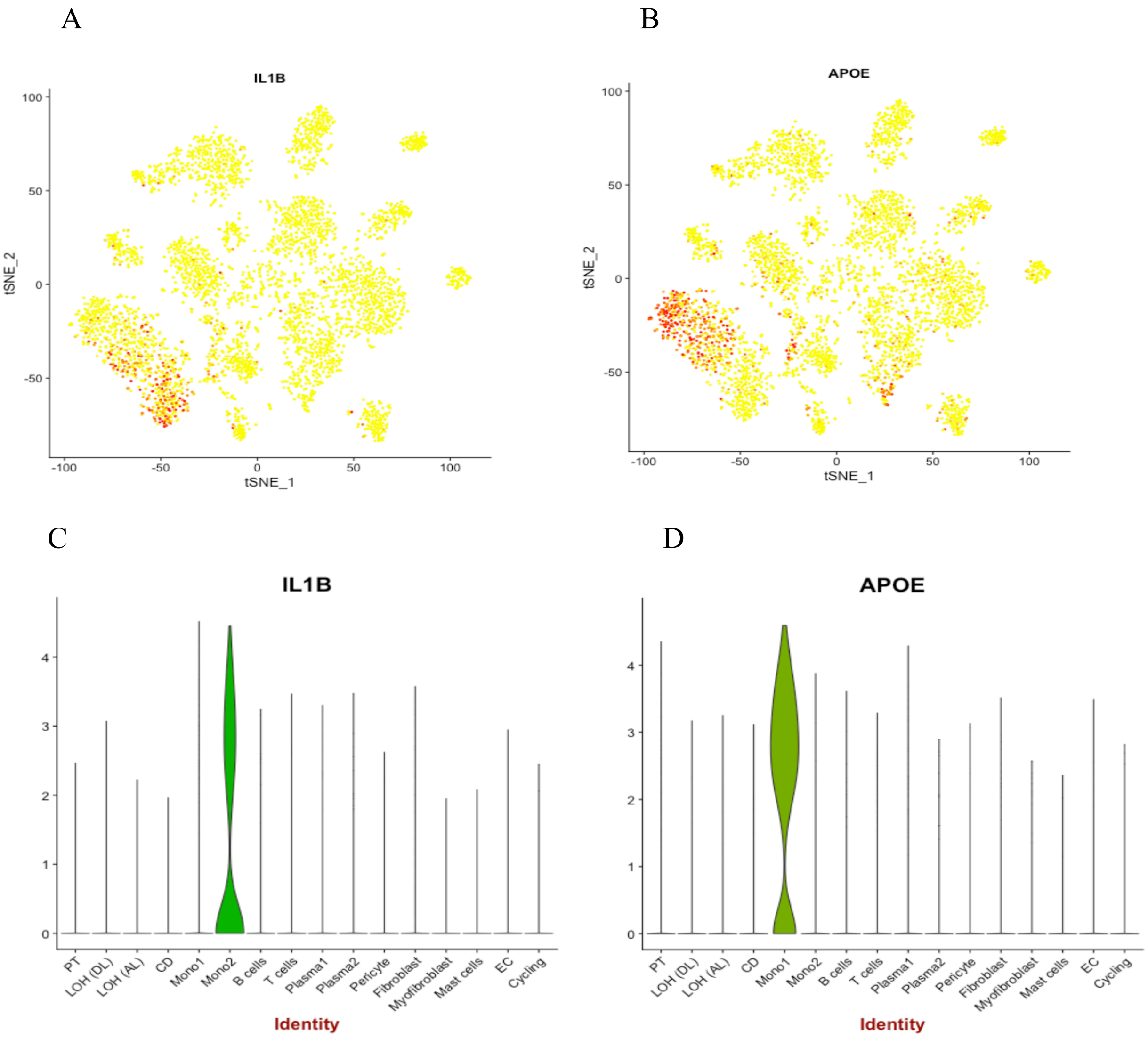


**Fig. S9.** The single cell RNA-seq data from human kidney allograft rejection from Wu and Malone et al, retrieved from Humphreys Lab. (**A**) t-SNE plot and (C) violin plot showing high *IL1B* expression in monocyte cluster 2 (Mono2). (**B**) t-SNE plot and (D) violin plot showing high *APOE* expression in monocyte cluster 1 (Mono1).

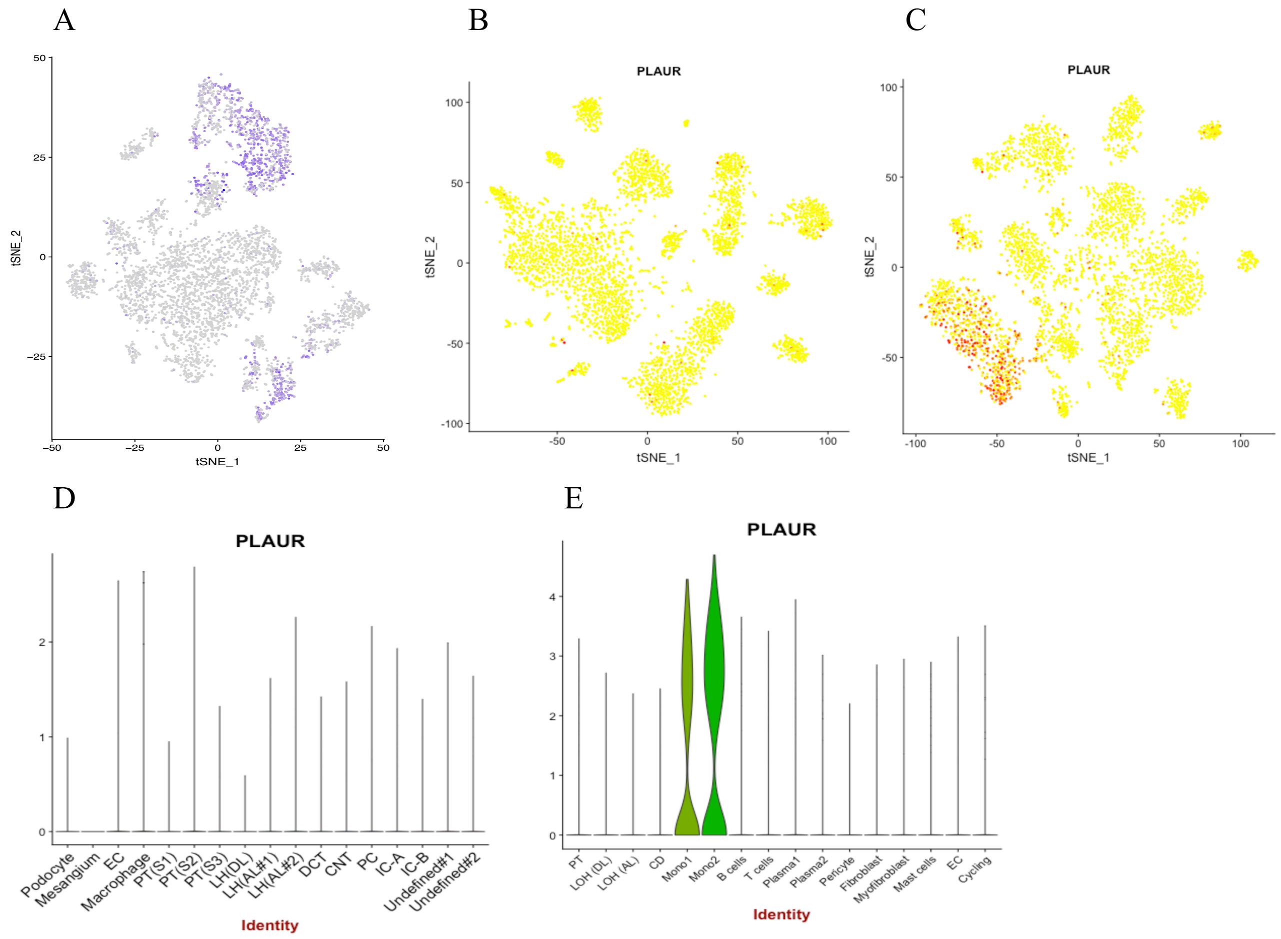


**Fig. S10.** High expression of *PLAUR* (encoding suPAR) found in monocyte clusters. (**A**) t-SNE plot of FSGS urine single cell data (current study). (**B**) t-SNE plot and (**D**) Violin plot of healthy human adult kidney single nucleus RNA-seq data from Wu and Uchimura et al. (**C**) t-SNE plot and (**E**) Violin plot of human rejecting kidney allograft data from Wu and Malone et al, retrieved from Humphreys Lab.

A

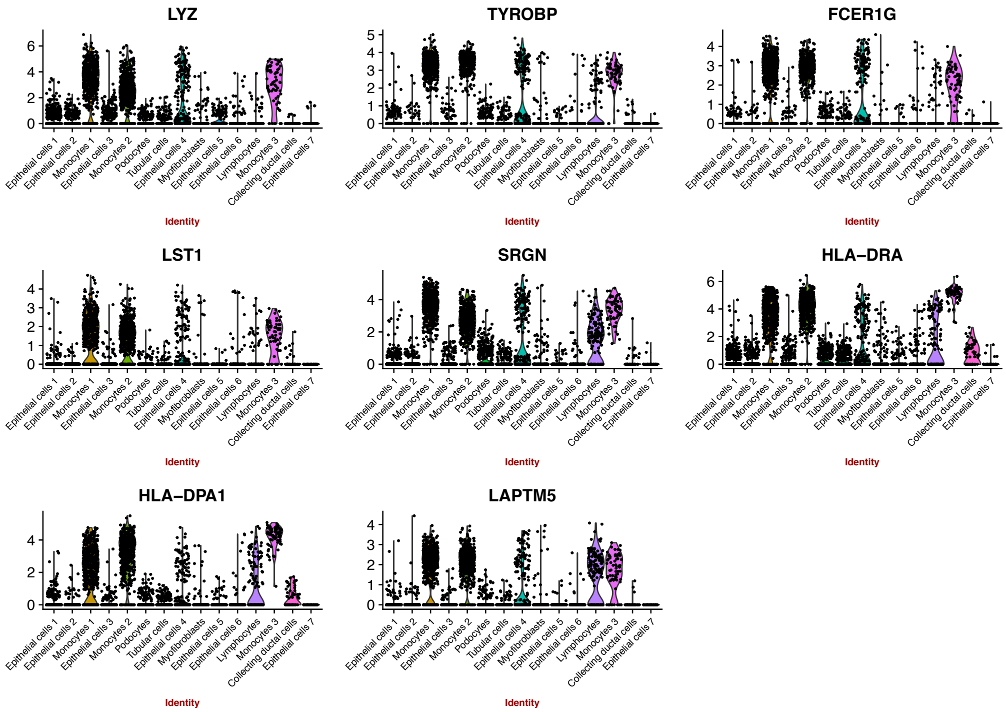


B

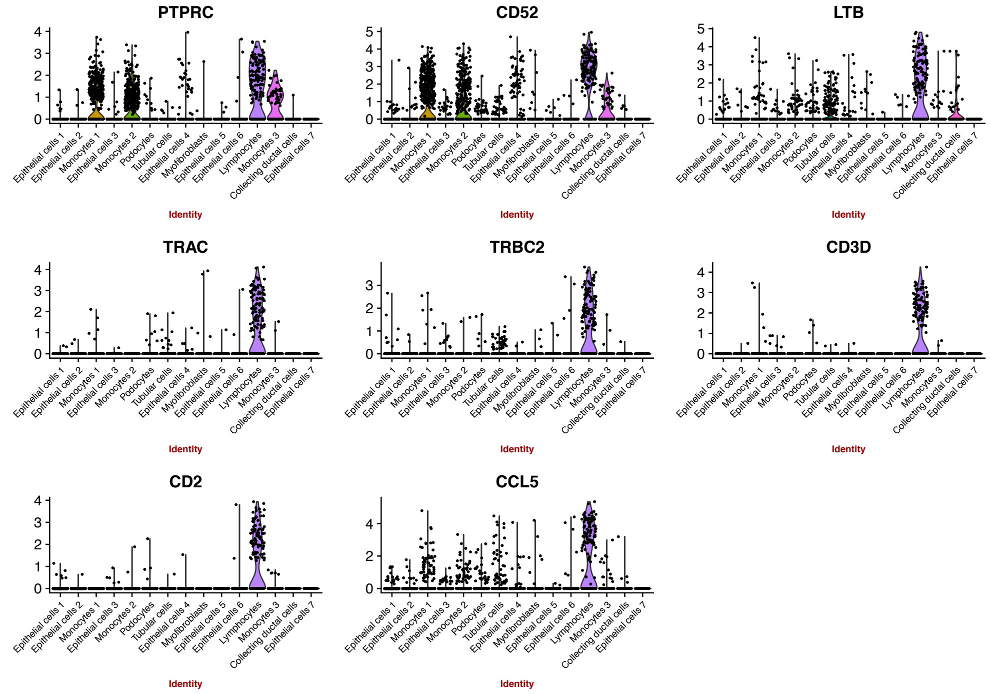


**Fig. S11.** Violin plots showing the most highly expressed monocyte/lymphocyte marker genes that were used to interrogate the presence of immune cells in the NEPTUNE transcriptomic data. (**A**) monocyte marker genes and (**B**) lymphocyte marker genes. Most of these genes showed little expression in other cell types.

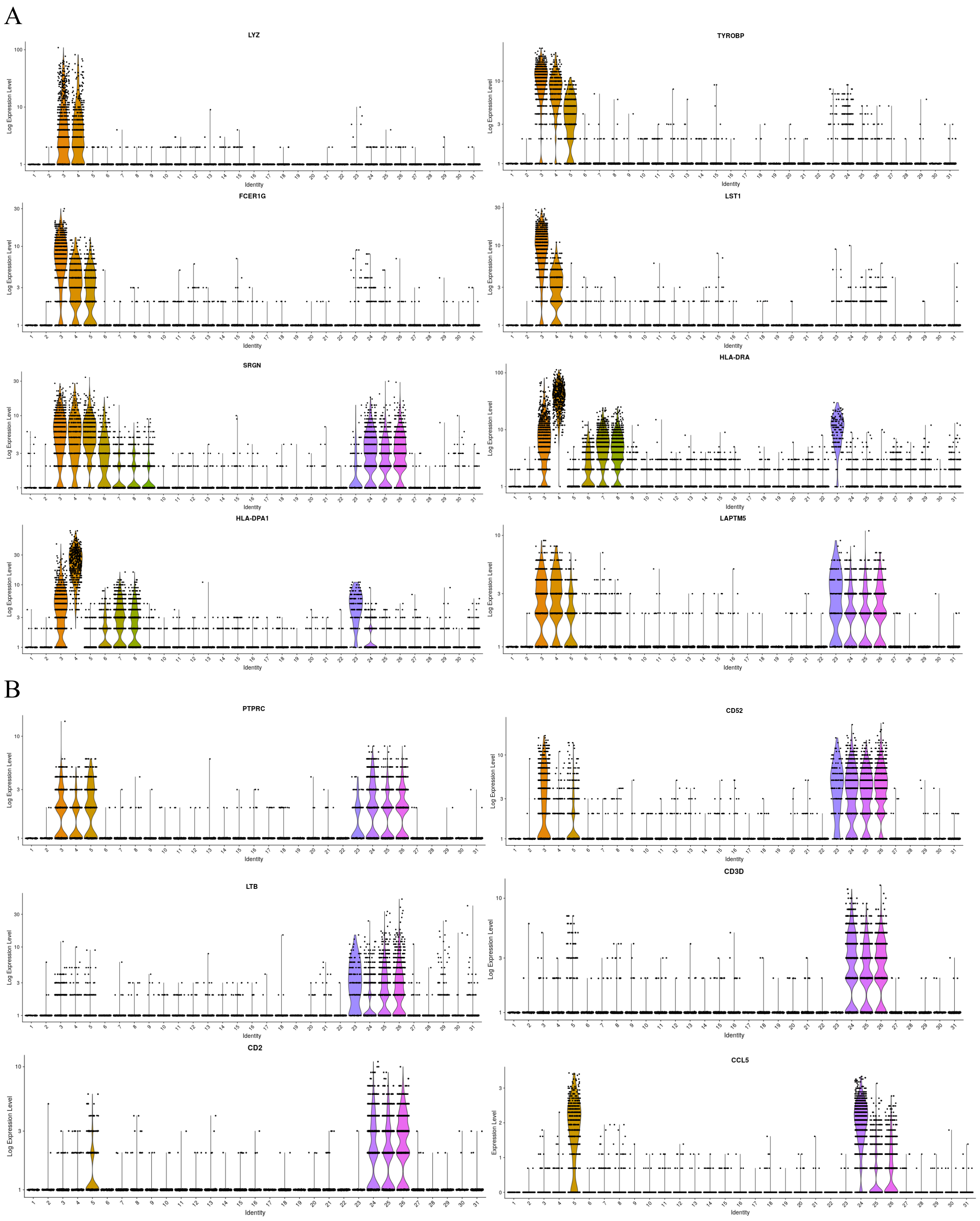


**Fig. S12.** Violin plots showing the most highly expressed monocyte/lymphocyte marker genes in the single nuclear RNAseq data from human adult kidney tissue from Menon et al. (**A**) monocyte marker genes (**B**) lymphocyte marker genes. (Cluster 3 = Macrophage; Cluster 4 = Monocyte; Cluster 5 = Natural killer cells; Cluster 6 = Endothelial cell (arteriolar); Cluster 7 = Endothelial cell (peritubular); Cluster 8 = Glomerular capillary endothelial cell; Cluster 9 = Vascular smooth muscle and mesangial cell; Cluster 23 = B cell; Cluster 24 = Cytotoxic T-cell; Cluster 25 = Activated T-cell; Cluster 26 = Memory T-cell). The expression levels of SRGN was ~3 times higher and the expression of *HLA-DRA* and *HLA-DPA1* genes was ~10 times higher in monocytes relative to endothelial cells. Two genes from urine lymphocyte cluster (*TRAC* and *TRBC2*) were not detected in this healthy kidney tissue snRNAseq dataset, possibly due to their specificity to T-cell activation state.

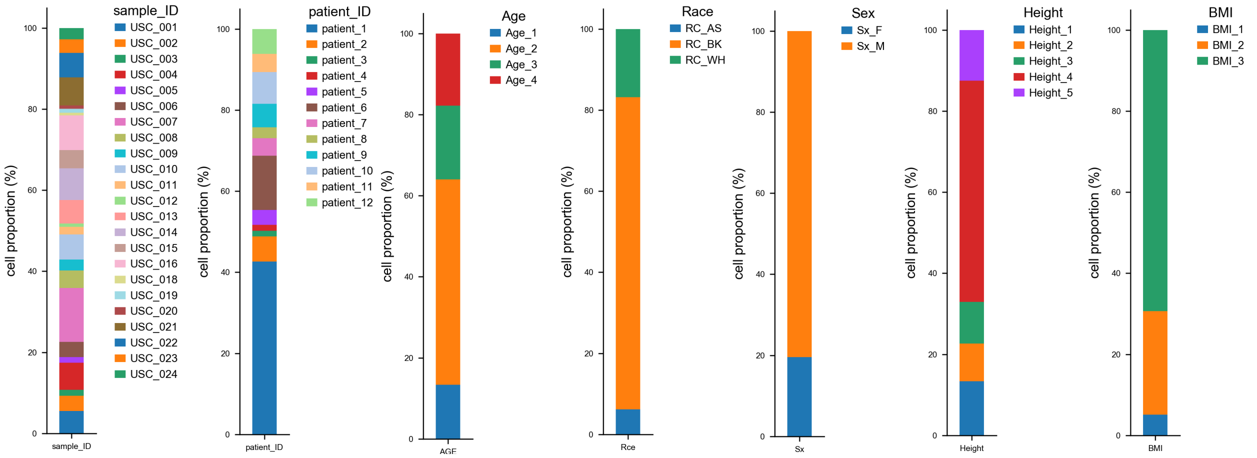


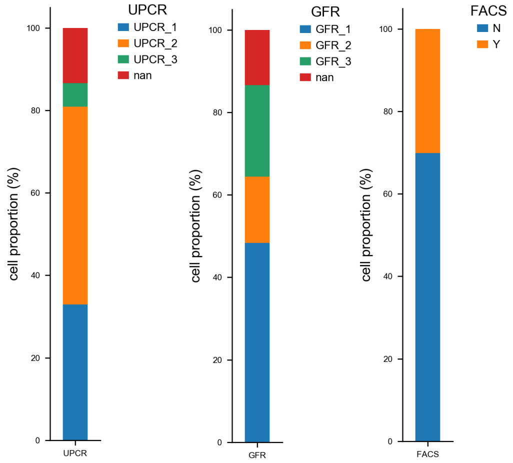


| Category | qualifier | N | bin_ID | | |
|---|---|---|---|---|---|
| Age | < 18 | 1 | Age_1 | | |
| | 19 - 40 | 4 | Age_2 | | |
| | 41 - 60 | 3 | Age_3 | | |
| | > 60 | 4 | Age_4 | | |
| Race | BK | 6 | RC_BK | | https://www.justice.gov/jm/eousa-resource-manual-70-racialethnic-codes |
| | WH | 5 | RC_WH | | |
| | AS | 1 | RC_AS | | |
| Sex | F | 5 | Sx_F | | |
| | M | 7 | Sx_M | | |
| GFR | 61 - 90 | 3 | GFR_1 | | |
| | 31 - 60 | 3 | GFR_2 | | |
| | 15 - 30 | 5 | GFR_3 | | |
| | NA | 1 | GFR_NA | | unknown GFR |
| UPCR | < 2 | 6 | UPCR_1 | | sub-nephrotic syndrome |
| | 2 - 5 | 4 | UPCR_2 | | nephrotic syndrome |
| | > 5 | 1 | UPCR_3 | | severe nephrotic syndrome (ns) |
| | NA | 1 | UPCR_NA | | unknown UPCR |
| BMI | 18.5 - 24.9 | 2 | BMI_1 | | normal |
| | 25 - 29.9 | 3 | BMI_2 | | overweight |
| | > 30 | 7 | BMI_3 | | obese |
| height | < 150 | 1 | Height_1 | | |
| | 150 - 159 | 2 | Height_2 | | |
| | 160 - 169 | 3 | Height_3 | | |
| | 170 - 180 | 3 | Height_4 | | |
| | >180 | 3 | Height_5 | | |
| no of samples | USC_001 | 7 | | | this donor provided 7 urine samples |
| | USC_002 | 4 | | | |
| | USC_011 | 2 | | | |
| | USC_023 | 2 | | | |
| | other 8 | 1 | | | these 8 donors provided 1 urine sample, each |
| diagnosis | FSGS | 9 | diag_FSGS | | |
| | CKD | 2 | diag_CKD | | |
| | MCD | 1 | diag_MCD | | |

**Fig. S13.** Stackplots showing cell population (%) by covariates. Sample ID, patient ID, age, race, sex, height, BMI, UPCR, GFR, and sample processing are shown. Table indicates the categorical keys.

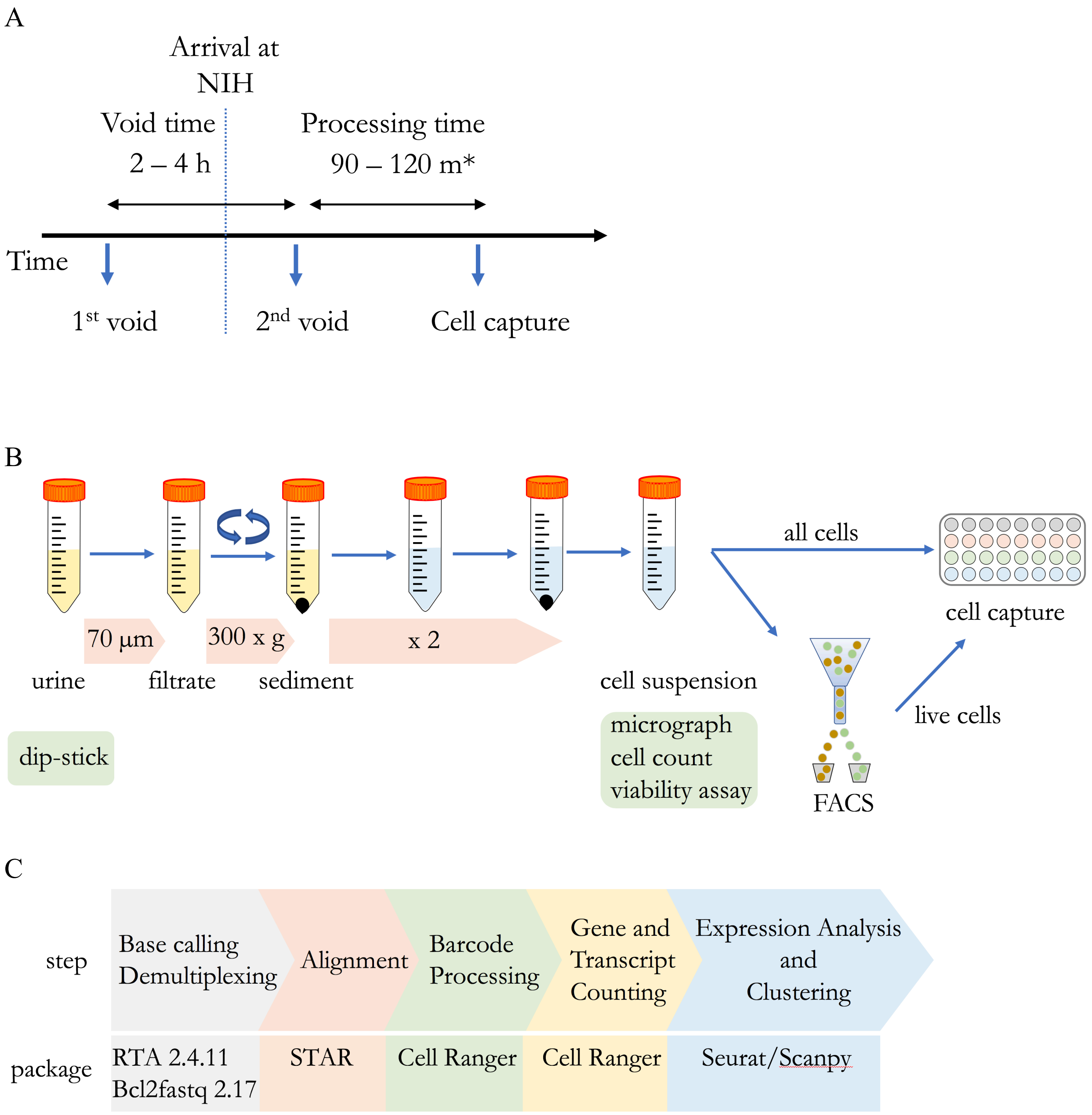


**Fig. S14.** Overview of the experiments and analysis of the urine single cell study. (**A**) Scheme for collecting urine and documenting time between 1st and 2nd void and processing time. *: this excludes a FACS step, which would add ~30 m to the processing time. (**B**) Scheme for urine processing and urine cell capture. (**C**) Key analysis steps and software utilized in the single-cell RNAseq data processing.

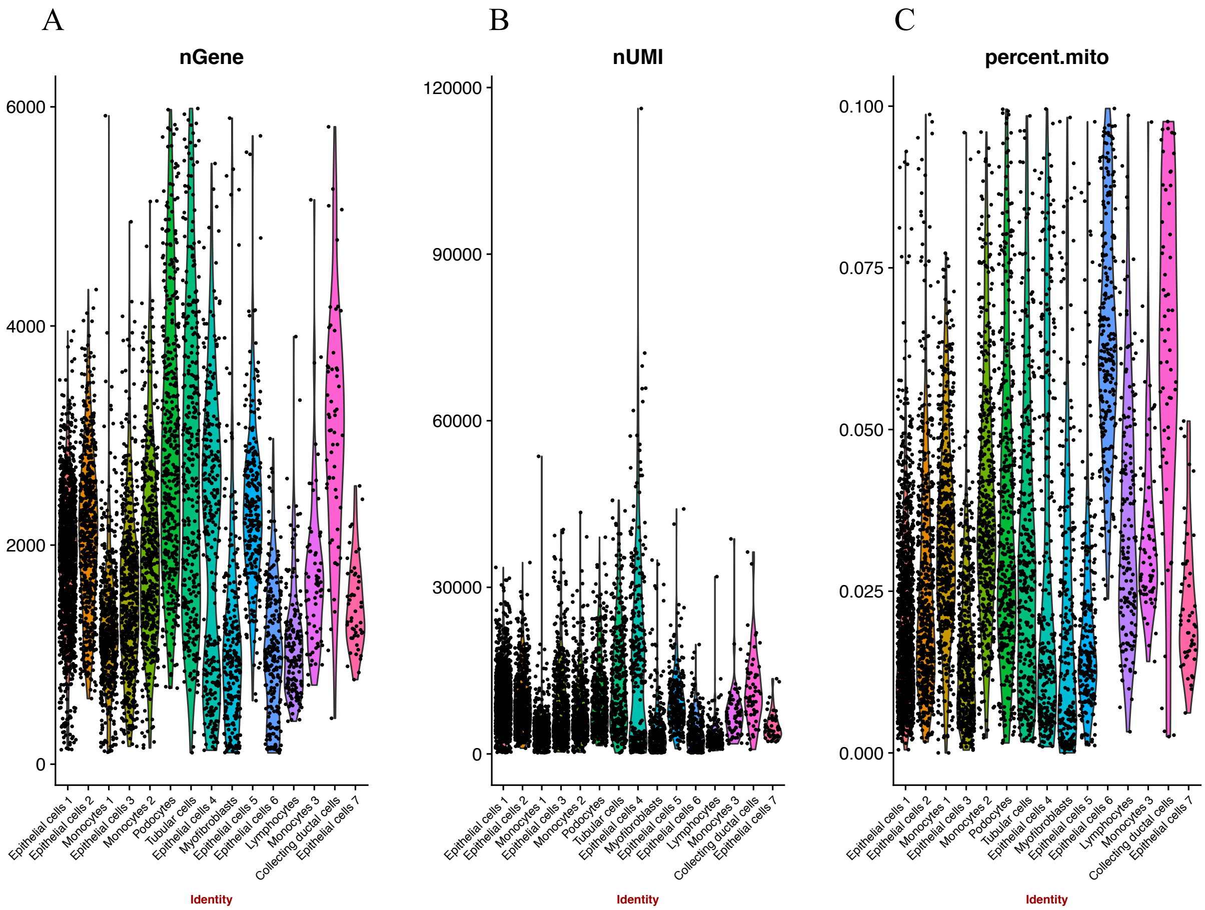


**Fig. S15.** Violin plots showing (**A**) number of genes (**B**) number of unique molecular identifiers and (**C**) mitochondrial percentage of individual clusters in the urine FSGS single cell dataset.

**Supplementary Tables:**

| Cell clusters | Subject 1 | Subject 2 | Subject 3 | Subject 4 | Subject 5 | Subject 6 | Subject 7 | Subject 8 | Subject 9 | Subject 10 | Subject 11 | Subject 12 |
|---|---|---|---|---|---|---|---|---|---|---|---|---|
| Epithelial cells 1 | 10.74 | 0.67 | 2.76 | 0.22 | 0.52 | 1.57 | 1.57 | 48.70 | 2.46 | 0.37 | 29.90 | 0.52 |
| Epithelial cells 2 | 10.14 | 0.52 | 1.75 | 0.35 | 0.17 | 1.92 | 2.97 | 31.99 | 20.80 | 0.17 | 26.57 | 2.62 |
| Epithelial cells 3 | 13.66 | 8.16 | 13.85 | 0.00 | 1.71 | 9.68 | 7.40 | 24.10 | 0.95 | 0.00 | 18.22 | 2.28 |
| Epithelial cells 4 | 22.97 | 12.72 | 5.65 | 0.71 | 3.18 | 0.00 | 1.06 | 20.14 | 2.12 | 11.31 | 18.02 | 2.12 |
| Epithelial cells 5 | 15.77 | 0.83 | 1.24 | 0.00 | 0.00 | 1.24 | 1.66 | 34.85 | 11.20 | 2.49 | 29.05 | 1.66 |
| Epithelial cells 6 | 16.81 | 6.03 | 17.67 | 1.29 | 3.88 | 1.29 | 5.17 | 12.93 | 1.72 | 0.43 | 28.88 | 3.88 |
| Epithelial cells 7 | 0.00 | 0.00 | 0.00 | 0.00 | 0.00 | 0.00 | 0.00 | 0.00 | 100.00 | 0.00 | 0.00 | 0.00 |
| Monocytes 1 | 39.37 | 12.13 | 1.76 | 0.70 | 4.39 | 0.00 | 1.05 | 0.70 | 0.00 | 36.73 | 0.88 | 2.28 |
| Monocytes 2 | 38.43 | 3.61 | 0.64 | 3.82 | 1.91 | 0.00 | 0.64 | 1.27 | 0.21 | 44.37 | 1.27 | 3.82 |
| Monocytes 3 | 22.39 | 5.97 | 0.00 | 2.99 | 4.48 | 0.00 | 4.48 | 1.49 | 1.49 | 41.79 | 10.45 | 4.48 |
| Lymphocytes | 41.72 | 13.50 | 1.23 | 9.20 | 4.29 | 0.00 | 1.84 | 1.84 | 0.00 | 4.29 | 19.02 | 3.07 |
| Podocytes | 56.64 | 4.51 | 0.00 | 5.76 | 0.00 | 0.50 | 0.75 | 4.76 | 5.51 | 16.04 | 4.01 | 1.50 |
| Tubular cells | 45.45 | 2.73 | 0.91 | 6.06 | 0.91 | 0.30 | 1.52 | 10.91 | 10.61 | 6.36 | 7.88 | 6.36 |
| Collecting ductal cells | 74.58 | 0.00 | 1.69 | 3.39 | 0.00 | 0.00 | 0.00 | 1.69 | 3.39 | 11.86 | 1.69 | 1.69 |
| Myofibroblasts | 39.27 | 7.29 | 7.29 | 5.26 | 5.26 | 1.62 | 1.62 | 10.93 | 1.21 | 6.07 | 12.15 | 2.02 |

**Table S1.** The percentage of cells from each subject contributing to different cell clusters in the FSGS urine scRNA-seq study.

| Genes | Average log FC | pct.1 | pct.2 | Adjusted P |
|---|---|---|---|---|
| IGFBP7 | 2.54868202 | 0.845 | 0.24 | 7.31E-216 |
| CTGF | 2.05369501 | 0.519 | 0.107 | 2.15E-125 |
| SST | 2.03334856 | 0.223 | 0.02 | 3.11E-98 |
| SPON2 | 1.75554335 | 0.672 | 0.037 | 0 |
| CRYAB | 1.75461245 | 0.847 | 0.427 | 1.53E-124 |
| TPM1 | 1.64414834 | 0.797 | 0.162 | 4.58E-232 |
| PTGDS | 1.55664899 | 0.281 | 0.018 | 1.83E-155 |
| BCAM | 1.554537 | 0.672 | 0.078 | 6.65E-280 |
| NUPR1 | 1.50732189 | 0.744 | 0.162 | 4.03E-195 |
| VCAM1 | 1.47266711 | 0.546 | 0.016 | 0 |
| VIM | 1.43068533 | 0.85 | 0.345 | 2.64E-126 |
| CLU | 1.37849947 | 0.729 | 0.211 | 1.44E-146 |
| IFITM3 | 1.30483555 | 0.822 | 0.333 | 1.04E-125 |
| IL32 | 1.29664124 | 0.782 | 0.186 | 3.92E-177 |
| ARHGAP29 | 1.27788118 | 0.737 | 0.06 | 0 |
| SPARC | 1.27641137 | 0.586 | 0.036 | 0 |
| OCIAD2 | 1.27275393 | 0.722 | 0.139 | 4.44E-201 |
| TNFRSF12A | 1.27045897 | 0.802 | 0.317 | 1.15E-126 |
| MGP | 1.25322373 | 0.358 | 0.064 | 1.20E-93 |
| CALD1 | 1.20448825 | 0.734 | 0.048 | 0 |
| MYL9 | 1.20062467 | 0.604 | 0.08 | 5.43E-223 |
| TSC22D1 | 1.19580281 | 0.825 | 0.412 | 6.33E-91 |
| CYP1B1 | 1.19304652 | 0.449 | 0.046 | 1.41E-191 |
| CD151 | 1.18358191 | 0.842 | 0.348 | 4.17E-143 |
| S100A13 | 1.1724924 | 0.81 | 0.261 | 8.73E-161 |
| TUBA1A | 1.16192021 | 0.682 | 0.2 | 3.23E-129 |
| APP | 1.14365223 | 0.817 | 0.324 | 2.73E-139 |
| CLDN3 | 1.13502508 | 0.679 | 0.103 | 6.29E-226 |
| IGFBP4 | 1.11522485 | 0.526 | 0.025 | 0 |
| CALM2 | 1.06905551 | 0.937 | 0.644 | 3.90E-104 |
| GPX3 | 1.06402591 | 0.622 | 0.207 | 6.44E-97 |
| AKAP12 | 1.04537247 | 0.581 | 0.041 | 0 |
| ITM2B | 1.03844359 | 0.98 | 0.888 | 1.04E-75 |
| PLA2G16 | 1.03407815 | 0.757 | 0.236 | 1.58E-151 |
| FHL1 | 1.02157605 | 0.521 | 0.022 | 0 |
| ERP27 | 1.0185899 | 0.371 | 0.077 | 2.10E-84 |
| CAV1 | 1.00408385 | 0.571 | 0.055 | 1.68E-261 |
| FXYD2 | 1.00301828 | 0.581 | 0.099 | 1.60E-159 |

**Table S2.** The most highly expressed genes in podocyte cluster compared with remaining clusters. Pct.1 = percentage of cells in the podocyte cluster expressing the gene, pct.2 = percentage of all the remaining cells expressing the gene.

| GO Term | Annotated | Significant | Expected | Fisher |
|---|---|---|---|---|
| regulation of amyloid fibril formation | 8 | 3 | 0.02 | 6.00E-07 |
| cell adhesion | 1347 | 14 | 3.06 | 2.80E-05 |
| regulation of cell communication by electrical coupling involved in cardiac conduction | 9 | 2 | 0.02 | 0.00018 |
| positive regulation of amyloid-beta formation | 10 | 2 | 0.02 | 0.00022 |
| negative regulation of protein homooligomerization | 10 | 2 | 0.02 | 0.00022 |
| regulation of cell growth | 402 | 6 | 0.91 | 0.00027 |
| regulation of cardiac muscle contraction | 73 | 3 | 0.17 | 0.00061 |
| response to oxygen-containing compound | 1526 | 15 | 3.47 | 0.00086 |
| cartilage condensation | 20 | 2 | 0.05 | 0.00093 |
| extracellular matrix organization | 341 | 5 | 0.78 | 0.00099 |
| wound healing | 525 | 6 | 1.19 | 0.0011 |
| cell morphogenesis | 965 | 8 | 2.2 | 0.00125 |
| regulation of heart rate | 94 | 3 | 0.21 | 0.00126 |
| response to lead ion | 24 | 2 | 0.05 | 0.00135 |
| muscle organ development | 370 | 5 | 0.84 | 0.00143 |
| positive regulation of collagen biosynthetic process | 25 | 2 | 0.06 | 0.00146 |
| regulation of cell migration | 787 | 7 | 1.79 | 0.00176 |
| positive regulation of intracellular signal transduction | 1024 | 8 | 2.33 | 0.00183 |
| positive regulation of peptidyl-threonine phosphorylation | 29 | 2 | 0.07 | 0.00197 |
| negative regulation of cell proliferation | 660 | 8 | 1.5 | 0.00206 |

**Table S3.** Gene ontology pathway analysis of the most highly expressed genes (log FC >= 1) from podocyte cluster (n = 38). The numbers of annotated genes and the number of significant genes in each pathway are shown.

| Genes | Average log FC | pct.1 | pct.2 | Adjusted P |
|---|---|---|---|---|
| APOE | 3.41494627 | 0.636 | 0.18 | 9.42E-276 |
| HLA-DRA | 3.09999481 | 0.954 | 0.232 | 0 |
| CD74 | 2.97502743 | 0.949 | 0.349 | 0 |
| HLA-DPA1 | 2.81083669 | 0.868 | 0.115 | 0 |
| TYROBP | 2.70696373 | 0.978 | 0.112 | 0 |
| HLA-DRB1 | 2.70405694 | 0.92 | 0.241 | 0 |
| HLA-DPB1 | 2.69608228 | 0.868 | 0.133 | 0 |
| APOC1 | 2.6787553 | 0.592 | 0.117 | 0 |
| C1QB | 2.66786304 | 0.377 | 0.026 | 3.26E-282 |
| LYZ | 2.61580778 | 0.922 | 0.221 | 0 |
| SRGN | 2.53298644 | 0.977 | 0.168 | 0 |
| C1QA | 2.52854833 | 0.407 | 0.027 | 0 |
| FCER1G | 2.4206035 | 0.949 | 0.09 | 0 |
| TIMP1 | 2.40968273 | 0.869 | 0.237 | 0 |
| HLA-DRB5 | 2.33703663 | 0.859 | 0.082 | 0 |
| SPP1 | 2.25542253 | 0.598 | 0.201 | 4.25E-181 |
| AIF1 | 2.22434742 | 0.9 | 0.072 | 0 |
| CCL3 | 2.19146016 | 0.607 | 0.117 | 0 |
| C1QC | 2.12852785 | 0.358 | 0.018 | 1.96E-291 |
| FTL | 2.08625108 | 0.998 | 0.981 | 0 |
| HLA-DQA1 | 2.05276599 | 0.713 | 0.05 | 0 |
| PSAP | 2.04824297 | 0.946 | 0.616 | 0 |
| RP11-1143G9.4 | 1.95907451 | 0.575 | 0.08 | 0 |
| FABP5 | 1.94777774 | 0.653 | 0.128 | 0 |
| MT1H | 1.90705125 | 0.205 | 0.118 | 2.45E-13 |
| HLA-DQB1 | 1.88428731 | 0.704 | 0.104 | 0 |
| LGALS1 | 1.83008206 | 0.934 | 0.254 | 0 |
| MT1X | 1.82721852 | 0.538 | 0.239 | 9.52E-110 |
| LAPTM5 | 1.81992681 | 0.879 | 0.084 | 0 |
| RGS1 | 1.78397027 | 0.445 | 0.04 | 0 |
| HLA-DMA | 1.75892155 | 0.748 | 0.101 | 0 |
| CCL2 | 1.72524582 | 0.341 | 0.089 | 6.76E-107 |
| MT1G | 1.68411374 | 0.408 | 0.252 | 4.51E-35 |
| CD14 | 1.67413643 | 0.602 | 0.086 | 0 |
| CD68 | 1.6693397 | 0.799 | 0.373 | 0 |
| LST1 | 1.64223589 | 0.783 | 0.059 | 0 |

**Table S4.** The most highly expressed genes in monocyte clusters compared with remaining clusters. Pct.1 = percentage of cells in the monocyte clusters expressing the gene, pct.2 = percentage of all the remaining cells expressing the gene.

| GO Term | Annotated | Significant | Expected | Fisher |
|---|---|---|---|---|
| SRP-dependent cotranslational protein targeting to membrane | 97 | 80 | 4.34 | < 1e-30 |
| nuclear-transcribed mRNA catabolic process, nonsense-mediated decay | 120 | 79 | 5.37 | < 1e-30 |
| translational initiation | 191 | 88 | 8.55 | < 1e-30 |
| neutrophil degranulation | 480 | 113 | 21.5 | < 1e-30 |
| cytoplasmic translation | 90 | 37 | 4.03 | 1.20E-23 |
| platelet degranulation | 125 | 26 | 5.6 | 4.10E-11 |
| cytokine-mediated signaling pathway | 716 | 91 | 32.07 | 6.50E-11 |
| antigen processing and presentation of exogenous peptide antigen | 96 | 22 | 4.3 | 1.80E-10 |
| ribosomal small subunit assembly | 15 | 9 | 0.67 | 2.70E-09 |
| interferon-gamma-mediated signaling pathway | 87 | 19 | 3.9 | 7.40E-09 |
| inflammatory response | 754 | 89 | 33.77 | 7.70E-09 |
| cellular response to lipopolysaccharide | 192 | 36 | 8.6 | 3.20E-08 |
| positive regulation of T cell proliferation | 93 | 18 | 4.17 | 1.30E-07 |
| cellular response to interferon-gamma | 172 | 36 | 7.7 | 1.30E-07 |
| MHC protein complex assembly | 5 | 5 | 0.22 | 1.80E-07 |
| respiratory burst | 29 | 10 | 1.3 | 2.80E-07 |
| ribosomal large subunit assembly | 30 | 10 | 1.34 | 4.10E-07 |
| T cell receptor signaling pathway | 143 | 24 | 6.4 | 5.70E-07 |
| lipopolysaccharide-mediated signaling pathway | 55 | 13 | 2.46 | 6.70E-07 |
| regulation of cell shape | 148 | 22 | 6.63 | 7.20E-07 |

**Table S5.** Gene ontology pathway analysis of significant genes from monocyte clusters (n = 817). The numbers of annotated genes and the number of significant genes in each pathway are shown.

| Genes | Average log FC | pct.1 | pct.2 | Adjusted P |
|---|---|---|---|---|
| GNLY | 3.06721981 | 0.252 | 0.009 | 3.71E-128 |
| CCL5 | 2.9720821 | 0.564 | 0.051 | 1.34E-158 |
| LTB | 2.56875894 | 0.638 | 0.058 | 4.93E-182 |
| GZMA | 2.40914171 | 0.497 | 0.004 | 0 |
| CD52 | 2.40587021 | 0.853 | 0.135 | 7.01E-164 |
| NKG7 | 2.3523265 | 0.429 | 0.016 | 4.39E-222 |
| KLRB1 | 2.26163849 | 0.368 | 0.004 | 0 |
| IL32 | 2.1652586 | 0.718 | 0.214 | 4.49E-71 |
| TRAC | 2.12135384 | 0.601 | 0.008 | 0 |
| CD3D | 2.06180348 | 0.528 | 0.004 | 0 |
| CD2 | 2.01112974 | 0.528 | 0.005 | 0 |
| CD69 | 1.98266306 | 0.429 | 0.021 | 6.15E-181 |
| TRBC2 | 1.89469489 | 0.577 | 0.016 | 0 |
| TRBC1 | 1.86480698 | 0.405 | 0.004 | 0 |
| CORO1A | 1.85116787 | 0.767 | 0.199 | 4.32E-93 |
| CTSW | 1.74471433 | 0.436 | 0.011 | 5.10E-279 |
| GZMB | 1.71333409 | 0.258 | 0.009 | 1.07E-127 |
| PTPRC | 1.62606258 | 0.712 | 0.116 | 1.35E-119 |
| CST7 | 1.58724428 | 0.429 | 0.034 | 1.96E-125 |
| RPS27 | 1.53801411 | 0.994 | 0.959 | 4.02E-71 |
| RPS29 | 1.53485279 | 0.969 | 0.923 | 5.64E-69 |
| RAC2 | 1.49084526 | 0.62 | 0.104 | 5.86E-102 |
| LIMD2 | 1.48287126 | 0.583 | 0.127 | 5.54E-73 |
| GIMAP7 | 1.45319041 | 0.423 | 0.014 | 8.60E-228 |
| GZMH | 1.43683772 | 0.239 | 0.002 | 1.28E-219 |
| EVL | 1.4357777 | 0.546 | 0.098 | 4.05E-82 |
| CD7 | 1.43206526 | 0.411 | 0.031 | 1.40E-127 |
| HCST | 1.42680839 | 0.693 | 0.192 | 5.62E-61 |
| CCL4 | 1.41856257 | 0.319 | 0.122 | 4.65E-11 |
| CD3E | 1.41606845 | 0.417 | 0.004 | 0 |
| RPL23A | 1.37235222 | 0.945 | 0.855 | 2.36E-58 |
| CXCR4 | 1.35732602 | 0.595 | 0.142 | 1.21E-58 |
| RPS4X | 1.32584911 | 0.963 | 0.757 | 1.19E-55 |
| RPS15A | 1.30202755 | 0.982 | 0.918 | 3.96E-66 |
| AC092580.4 | 1.29063944 | 0.301 | 0.002 | 2.20E-290 |
| GZMK | 1.2808903 | 0.147 | 0.001 | 2.74E-125 |

**Table S6.** The most highly expressed genes in lymphocyte cluster compared with remaining clusters. Pct.1 = percentage of cells in the lymphocyte cluster expressing the gene, pct.2 = percentage of all the remaining cells expressing the gene.

| GO Term | Annotated | Significant | Expected | Fisher |
|---|---|---|---|---|
| SRP-dependent cotranslational protein targeting to membrane | 97 | 78 | 2.46 | < 1e-30 |
| nuclear-transcribed mRNA catabolic process, nonsense-mediated decay | 120 | 79 | 3.05 | < 1e-30 |
| translational initiation | 191 | 84 | 4.85 | < 1e-30 |
| cytoplasmic translation | 90 | 34 | 2.28 | < 1e-30 |
| ribosomal small subunit assembly | 15 | 9 | 0.38 | 1.80E-11 |
| ribosomal large subunit assembly | 30 | 11 | 0.76 | 8.80E-11 |
| T cell receptor signaling pathway | 143 | 25 | 3.63 | 9.60E-11 |
| adaptive immune response | 596 | 59 | 15.13 | 8.90E-10 |
| immune response | 2112 | 160 | 53.62 | 1.80E-09 |
| positive thymic T cell selection | 13 | 7 | 0.33 | 9.70E-09 |
| cellular defense response | 57 | 12 | 1.45 | 1.50E-08 |
| positive regulation of T cell proliferation | 93 | 18 | 2.36 | 3.40E-08 |
| rRNA processing | 192 | 28 | 4.87 | 7.00E-08 |
| negative thymic T cell selection | 11 | 6 | 0.28 | 1.10E-07 |
| B cell receptor signaling pathway | 110 | 17 | 2.79 | 2.20E-07 |
| regulation of immune response | 977 | 91 | 24.8 | 2.60E-07 |
| positive regulation of T cell mediated cytotoxicity | 24 | 7 | 0.61 | 1.50E-06 |
| Fc-gamma receptor signaling pathway involved in phagocytosis | 134 | 15 | 3.4 | 1.60E-06 |
| interleukin-12-mediated signaling pathway | 47 | 9 | 1.19 | 2.30E-06 |
| regulation of defense response to virus by virus | 29 | 7 | 0.74 | 6.20E-06 |

**Table S7.** Gene ontology pathway analysis of significant genes from lymphocyte cluster (n = 481). The numbers of annotated genes and the number of significant genes in each pathway are shown.

| Gene | Fold change | Mean Expression of low subset | Mean Expression of high subset | Adj. p-value |
|---|---|---|---|---|
| RNASE1 | 3.0841 | 0.7077 | 2.3909 | 1.25E-05 |
| A2M* | 2.9296 | 0.6975 | 2.2362 | 1.48E-07 |
| SEPP1* | 2.6828 | 0.8243 | 2.3797 | 1.67E-04 |
| GPNMB* | 2.6802 | 0.8291 | 2.3902 | 3.13E-07 |
| C1QC* | 2.5765 | 0.8077 | 2.2387 | 1.48E-07 |
| C1QB* | 2.4191 | 1.0703 | 2.731 | 6.62E-07 |
| LGMN* | 2.4101 | 1.0296 | 2.6225 | 3.45E-07 |
| C1QA* | 2.3089 | 0.9311 | 2.2806 | 4.20E-06 |
| SPP1* | 2.2675 | 1.2821 | 3.034 | 4.95E-04 |
| VSIG4* | 2.1473 | 0.9457 | 2.1454 | 3.17E-05 |
| CTSL1 | 1.7209 | 1.2786 | 2.2724 | 1.75E-03 |
| CTSD | 1.6807 | 2.322 | 3.9706 | 1.91E-08 |
| LIPA | 1.6772 | 1.26 | 2.181 | 7.02E-04 |
| ASAH1 | 1.5316 | 1.4947 | 2.3424 | 1.46E-04 |
| FCGR3A | 1.5134 | 1.6875 | 2.6052 | 1.79E-02 |
| CD14 | 1.5062 | 2.0646 | 3.1605 | 1.50E-03 |
| FCGRT | 1.4808 | 1.5147 | 2.291 | 1.59E-03 |
| CD163 | 1.4724 | 1.372 | 2.0674 | 2.44E-02 |
| FCGR2A | 1.3958 | 1.5922 | 2.262 | 1.09E-02 |
| CD63 | 1.3595 | 1.5918 | 2.2 | 4.87E-03 |
| CSF1R | 1.3562 | 1.8538 | 2.5498 | 3.75E-02 |
| CD68* | 1.2797 | 2.4132 | 3.1163 | 1.94E-04 |
| CTSB | 1.2627 | 3.695 | 4.692 | 5.60E-05 |
| GPX1* | 1.2545 | 1.9061 | 2.4166 | 6.50E-03 |
| NPC2* | 1.2061 | 1.9794 | 2.408 | 2.37E-02 |

**Table S8.** Top 25 upregulated genes in *APOE*$^{+}$ macrophages when compared with *APOE*$^{-}$ macrophages in untreated melanoma from Jerby-Arnon et al. Marked with asterisk are the genes that were also most highly expressed in M2 monocytes from FSGS urine single cell data.

| Gene | Fold change | Mean Expression of low subset | Mean Expression of high subset | Adj. p-value |
|---|---|---|---|---|
| FABP3 | 7.1922 | 0.2321 | 2.2883 | 1.33E-04 |
| ACE | 7.145 | 0.2133 | 2.1384 | 1.60E-06 |
| APOC1* | 4.6155 | 0.9554 | 4.7713 | 4.52E-06 |
| NUPR1 | 4.1999 | 0.6517 | 3.0572 | 1.18E-03 |
| VMO1 | 4.0273 | 0.4993 | 2.3137 | 1.17E-02 |
| SLC29A1 | 3.9456 | 0.4752 | 2.1697 | 1.85E-03 |
| GPNMB* | 3.7242 | 1.6418 | 6.3867 | 4.47E-07 |
| TMEM37* | 3.664 | 0.5951 | 2.4467 | 1.32E-03 |
| PLA2G7 | 3.5473 | 1 | 3.8019 | 1.33E-04 |
| FUCA1 | 3.4359 | 0.9074 | 3.3612 | 9.25E-04 |
| CD209 | 3.4114 | 0.6224 | 2.3643 | 1.70E-02 |
| SLC1A3 | 3.3712 | 0.852 | 3.1093 | 2.57E-04 |
| C2 | 3.2867 | 1.2627 | 4.3787 | 1.33E-04 |
| HGF | 3.2268 | 0.5641 | 2.0428 | 1.64E-02 |
| RAB13 | 3.1583 | 0.7804 | 2.6806 | 5.44E-03 |
| C1orf85 | 3.0873 | 1.0637 | 3.4927 | 3.16E-04 |
| GGA1 | 3.0568 | 0.6669 | 2.2441 | 1.87E-02 |
| KDELR1 | 3.0262 | 0.649 | 2.1665 | 7.92E-03 |
| TREM2* | 3.0063 | 1.3981 | 4.4037 | 9.17E-05 |
| A2M* | 2.9218 | 1.7123 | 5.195 | 5.33E-05 |
| SPP1* | 2.8944 | 2.0877 | 6.2322 | 7.66E-03 |
| SLC38A6 | 2.8841 | 0.6902 | 2.179 | 2.35E-02 |
| SLC7A8 | 2.8205 | 1.013 | 3.0392 | 1.09E-03 |
| PMP22 | 2.7991 | 1.3969 | 4.0901 | 4.07E-04 |
| ACP5 | 2.7816 | 1.8829 | 5.4156 | 1.54E-05 |

**Table S9.** Top 25 upregulated genes in *APOE*$^{+}$ macrophages when compared with *APOE*$^{-}$ macrophages in head and neck cancer from Puram et al. Marked with asterisk are the genes that were also most highly expressed in M2 monocytes from FSGS urine single cell data.

| Gene | Fold change | Mean Expression of low subset | Mean Expression of high subset | Adj. p-value |
|---|---|---|---|---|
| TNFAIP3 | 3.5537 | 0.5307 | 2.1414 | 5.81E-12 |
| SOD2* | 3.0435 | 0.6155 | 2.0777 | 1.79E-10 |
| NAMPT* | 2.9996 | 0.6385 | 2.1153 | 3.08E-11 |
| PPP1R15A | 2.7635 | 0.8061 | 2.4041 | 7.82E-11 |
| NFKBIA | 2.679 | 0.9816 | 2.7977 | 6.47E-13 |
| JUNB | 2.6635 | 0.8708 | 2.4858 | 3.08E-11 |
| DUSP1 | 2.1342 | 1.7919 | 3.9378 | 1.01E-13 |
| ZFP36 | 2.1191 | 1.2134 | 2.6833 | 3.47E-10 |
| SERPINA1 | 1.8733 | 1.194 | 2.3241 | 1.03E-05 |
| CD44* | 1.7589 | 1.1085 | 2.0255 | 1.91E-06 |
| FOS | 1.6682 | 2.2153 | 3.7623 | 6.31E-11 |
| EIF1 | 1.5757 | 1.4359 | 2.3202 | 1.04E-10 |
| KLF6 | 1.5445 | 1.308 | 2.0746 | 3.10E-05 |
| LYZ | 1.5071 | 2.546 | 3.8877 | 4.87E-05 |
| PTPRC | 1.413 | 1.4729 | 2.1225 | 5.86E-04 |
| ZFP36L1 | 1.3691 | 1.4756 | 2.0571 | 2.63E-02 |
| LCP1 | 1.2632 | 1.8948 | 2.4199 | 2.05E-02 |
| TAPBP | 1.2373 | 1.7024 | 2.13 | 2.58E-02 |
| SH3BGRL3 | 1.2366 | 1.8427 | 2.3023 | 9.59E-03 |
| SRGN* | 1.2161 | 2.4708 | 3.0265 | 1.93E-03 |
| DDX5 | 1.2 | 2.1309 | 2.577 | 1.04E-02 |
| PSAP | 0.9187 | 4.436 | 4.0672 | 2.82E-02 |
| FTL | 0.8667 | 4.997 | 4.3174 | 6.11E-04 |
| PPIA | 0.8614 | 2.3537 | 2.0137 | 2.74E-03 |
| GRN | 0.8521 | 3.4543 | 2.9287 | 1.62E-02 |

**Table S10.** Top 25 upregulated genes in *IL1B*$^{+}$ macrophages when compared with *IL1B*$^{-}$ macrophages in untreated melanoma from Jerby-Arnon et al. Marked with asterisk are the genes that were also most highly expressed in M1 monocytes from FSGS urine single cell data.

| Gene | Fold change | Mean Expression of low subset | Mean Expression of high subset | Adj. p-value |
|---|---|---|---|---|
| LGALS2 | 6.9602 | 0.2667 | 2.4523 | 4.10E-03 |
| PMAIP1 | 5.3271 | 0.4975 | 3.0828 | 3.59E-03 |
| TNF | 3.4074 | 0.7079 | 2.6528 | 2.84E-02 |
| IL8* | 2.7916 | 1.8461 | 5.3327 | 2.84E-02 |
| IER3 | 2.6893 | 1.1086 | 3.1504 | 2.06E-02 |
| DUSP2 | 2.2697 | 2.0841 | 4.8573 | 2.57E-02 |
| TNFAIP3 | 2.1037 | 2.3173 | 4.9852 | 1.13E-02 |
| NFKBIZ | 2.0973 | 1.9058 | 4.1068 | 2.56E-02 |
| BCL2A1* | 1.8422 | 3.0902 | 5.7769 | 1.13E-02 |
| SERPINB9* | 1.7359 | 2.9757 | 5.2392 | 1.08E-02 |
| NR4A1 | 1.6848 | 3.2683 | 5.575 | 4.43E-02 |
| IER2 | 1.6557 | 3.4269 | 5.7393 | 2.06E-02 |
| PPP1R15A | 1.6261 | 3.8875 | 6.384 | 1.08E-02 |
| CD83 | 1.4648 | 4.2903 | 6.331 | 2.06E-02 |
| NFKBIA | 1.4198 | 5.7145 | 8.1553 | 1.27E-02 |
| GPNMB | 0.3367 | 4.3552 | 1.4002 | 2.57E-02 |
| DHRS3 | 0.2003 | 2.8839 | 0.4976 | 1.79E-02 |

**Table S11.** Top 17 upregulated genes in *IL1B*$^{+}$ macrophages when compared with *IL1B*$^{-}$ macrophages in head and neck cancer from Puram et al. Marked with asterisk are the genes that were also most highly expressed in M1 monocytes from FSGS urine single cell data.

| | Genes | Average log FC | pct.1 | pct.2 | Adjusted P |
|---|---|---|---|---|---|
| **Monocyte cluster genes** | HLA-DRA | 3.09999481 | 0.954 | 0.232 | 0 |
| | HLA-DPA1 | 2.81083669 | 0.868 | 0.115 | 0 |
| | TYROBP | 2.70696373 | 0.978 | 0.112 | 0 |
| | LYZ | 2.61580778 | 0.922 | 0.221 | 0 |
| | SRGN | 2.53298644 | 0.977 | 0.168 | 0 |
| | FCER1G | 2.4206035 | 0.949 | 0.09 | 0 |
| | LAPTM5 | 1.81992681 | 0.879 | 0.084 | 0 |
| | LST1 | 1.64223589 | 0.783 | 0.059 | 0 |
| **Lymphocyte cluster genes** | CCL5 | 2.9720821 | 0.564 | 0.051 | 1.34E-158 |
| | LTB | 2.56875894 | 0.638 | 0.058 | 4.93E-182 |
| | CD52 | 2.40587021 | 0.853 | 0.135 | 7.01E-164 |
| | TRAC | 2.12135384 | 0.601 | 0.008 | 0 |
| | CD3D | 2.06180348 | 0.528 | 0.004 | 0 |
| | CD2 | 2.01112974 | 0.528 | 0.005 | 0 |
| | TRBC2 | 1.89469489 | 0.577 | 0.016 | 0 |
| | PTPRC | 1.62606258 | 0.712 | 0.116 | 1.35E-119 |

**Table S12.** The 16 most highly expressed genes from monocyte and lymphocyte clusters selected to be evaluated in the NEPTUNE kidney transcriptomic data. They were selected based on their high levels of expression in their own clusters and relatively low levels of expression in the remaining cells. Pct.1 = percentage of cells in the clusters expressing the gene, pct.2 = percentage of all the remaining cells expressing the gene.

| Genes | Average log FC | pct.1 | pct.2 | Adjusted P |
|---|---|---|---|---|
| CTGF | 2.05369501 | 0.519 | 0.107 | 2.15E-125 |
| CRYAB | 1.75461245 | 0.847 | 0.427 | 1.53E-124 |
| TPM1 | 1.64414834 | 0.797 | 0.162 | 4.58E-232 |
| VIM | 1.43068533 | 0.85 | 0.345 | 2.64E-126 |
| CALD1 | 1.20448825 | 0.734 | 0.048 | 0 |
| MYL9 | 1.20062467 | 0.604 | 0.08 | 5.43E-223 |
| CD151 | 1.18358191 | 0.842 | 0.348 | 4.17E-143 |
| CAV1 | 1.00408385 | 0.571 | 0.055 | 1.68E-261 |
| TAGLN | 0.97754759 | 0.361 | 0.056 | 1.13E-103 |
| THY1 | 0.94158089 | 0.429 | 0.019 | 9.49E-290 |

**Table S13.** Ten genes from the podocyte cluster reported to be associated with EMT. Pct.1 = percentage of cells in the podocyte cluster expressing the gene, pct.2 = percentage of all the remaining cells expressing the gene.

| Study participants | Age | Sex | Ethnicity | Urine protein to creatinine ratio | eGFR | Steroid responsiveness | APOL1 status |
|---|---|---|---|---|---|---|---|
| Subject 1 (7 samples) | 39 | Male | African American | 4.23 | 88 | Resistant | G1G2 |
| Subject 2 (4 samples) | 47 | Female | African American | 3.18 | 38 | Partial remission | Not available |
| Subject 3 (2 samples) | 70 | Female | African American | 0.22 | 28 | Resistant | G1G0 |
| Subject 4 (2 samples) | 38 | Male | Asian | 2.83 | 45 | Not on steroid | G0G0 |
| Subject 5 (1 sample) | 62 | Female | European American | 4.84 | 25 | Resistant | G0G0 |
| Subject 6 (1 sample) | 50 | Female | European American | 0.39 | 85 | Not on steroid | Not available |
| Subject 7 (1 sample) | 29 | Male | European American | 0.17 | 54 | Not on steroid | Not available |
| Subject 8 (1 sample) | 15 | Male | African American | Not available | Not available | Partial remission | Not available |
| Subject 9 (1 sample) | 19 | Male | African American | Undetectable | 79 | Not on steroid | Not available |
| Subject 10 (1 sample) | 76 | Male | European American | 11.06 | 17 | Resistant | G0G0 |
| Subject 11 (1 sample) | 65 | Female | African American | 0.77 | 30 | Resistant | G0G0 |
| Subject 12 (1 sample) | 53 | Male | European American | 0.62 | 18 | Not on steroid | Not available |

**Table S14.** Demographic characteristics of participants in the FSGS urine single cell RNA-seq study.

| Sample | Subject | Estimated Number of Cells | Number of Reads | Mean Reads per Cell | Valid Barcodes | Total Genes Detected | Median Genes per Cell | Median UMI Counts per Cell |
|---|---|---|---|---|---|---|---|---|
| Sample_001 | Subject 1 | 303 | 96,282,471 | 317,763 | 97.60% | 19,957 | 2,513 | 11,889 |
| Sample_002 | Subject 2 | 105 | 138,291,330 | 1,317,060 | 97.40% | 13,375 | 1,020 | 3,545 |
| Sample_003 | Subject 5 | 135 | 108,821,417 | 806,084 | 96.10% | 12,969 | 203 | 452 |
| Sample_004 | Subject 1 | 362 | 120,133,413 | 331,860 | 95.80% | 19,363 | 1,003 | 3,589 |
| Sample_005 | Subject 6 | 125 | 128,858,236 | 1,030,865 | 84.30% | 10,887 | 670 | 2,017 |
| Sample_006 | Subject 7 | 173 | 118,805,305 | 686,735 | 91.90% | 15,514 | 1,243 | 4,525 |
| Sample_007 | Subject 8 | 1,264 | 114,163,025 | 90,318 | 97.10% | 18,691 | 1,895 | 8,847 |
| Sample_008 | Subject 9 | 317 | 69,035,478 | 217,777 | 97.90% | 15,955 | 1,880 | 6,844 |
| Sample_009 | Subject 1 | 726 | 52,884,441 | 72,843 | 97.50% | 15,195 | 71 | 1,263 |
| Sample_010 | Subject 1 | 244 | 51,839,761 | 212,458 | 97.50% | 17,475 | 2,047 | 10,532 |
| Sample_011 | Subject 3 | 174 | 73,343,018 | 421,511 | 97.00% | 11,147 | 199 | 526 |
| Sample_012 | Subject 3 | 105 | 60,066,783 | 572,064 | 97.90% | 12,651 | 1,240 | 4,897 |
| Sample_013 | Subject 10 | 778 | 66,151,706 | 85,027 | 97.60% | 20,038 | 1,573 | 6,097 |
| Sample_014 | Subject 11 | 994 | 53,378,029 | 53,700 | 97.40% | 18,648 | 1,891 | 8,296 |
| Sample_015 | Subject 12 | 391 | 50,210,220 | 128,414 | 97.00% | 16,568 | 181 | 1,476 |
| Sample_016 | Subject 1 | 364 | 72,612,789 | 199,485 | 97.80% | 18,321 | 1,436 | 4,078 |
| Sample_018 | Subject 2 | 756 | 68,247,170 | 90,274 | 98.00% | 10,451 | 38 | 294 |
| Sample_019 | Subject 2 | 239 | 66,384,275 | 277,758 | 98.10% | 12,070 | 90 | 390 |
| Sample_020 | Subject 2 | 519 | 61,667,357 | 118,819 | 97.70% | 12,511 | 39 | 338 |
| Sample_021 | Subject 1 | 384 | 105,479,299 | 274,685 | 97.10% | 18,309 | 959 | 2,988 |
| Sample_022 | Subject 1 | 519 | 116,170,509 | 223,835 | 97.30% | 18,370 | 574 | 2,443 |
| Sample_023 | Subject 4 | 1,559 | 122,004,786 | 78,258 | 97.00% | 16,697 | 48 | 1,187 |
| Sample_024 | Subject 4 | 1,938 | 112,444,649 | 58,020 | 97.40% | 16,595 | 38 | 956 |

**Table S15.** Statistics of the cell counts, barcodes, reads, genes and UMIs for each sample.
